# Chromosome 7 to the rescue: overcoming chromosome 10 loss in gliomas

**DOI:** 10.1101/2024.01.17.576103

**Authors:** Nishanth Ulhas Nair, Alejandro A. Schäffer, E. Michael Gertz, Kuoyuan Cheng, Johanna Zerbib, Avinash Das Sahu, Gil Leor, Eldad D. Shulman, Kenneth D. Aldape, Uri Ben-David, Eytan Ruppin

## Abstract

The co-occurrence of chromosome 10 loss and chromosome 7 gain in gliomas is the most frequent loss-gain co-aneuploidy pair in human cancers, a phenomenon that has been investigated without resolution since the late 1980s. Expanding beyond previous gene-centric studies, we investigate the co-occurrence in a genome-wide manner taking an evolutionary perspective. First, by mining large tumor aneuploidy data, we predict that the more likely order is 10 loss followed by 7 gain. Second, by analyzing extensive genomic and transcriptomic data from both patients and cell lines, we find that this co-occurrence can be explained by functional rescue interactions that are highly enriched on 7, which can possibly compensate for any detrimental consequences arising from the loss of 10. Finally, by analyzing transcriptomic data from normal, non-cancerous, human brain tissues, we provide a plausible reason why this co-occurrence happens preferentially in cancers originating in certain regions of the brain.

## INTRODUCTION

Gliomas, including glioblastoma (GBM) and lower-grade gliomas (LGG), present challenges for treatment due to their poor prognosis^1^. Since the late 1980’s it has been known that loss of chromosome 10 (10 loss) and gain of chromosome 7 (7 gain) often co-occur in GBM^2–5^. These two copy number alterations (CNAs) occur predominantly in IDH-wildtype GBM^6,7^, which compared to the *IDH1/IDH2*-mutant form has a worse prognosis^8^. Within the IDH-wildtype GBM category, the presence of 10 loss/7 gain was not significantly associated with prognosis in a multivariate clustering approach^9^, but segmental loss of intervals within chromosome 10 was associated with a better prognosis than the loss of the entire chromosome 10^10^. Previous studies aiming to order the two CNAs suggested that both 10 loss and 7 gain are early events in tumorigenesis that can occur in either order, but with 10 loss more commonly preceding 7 gain^3,6,7,11,12^.

Genomic studies in 1990’s and 2000’s used techniques such as microsatellite marker genotyping and comparative genomic hybridization (CGH) to characterize segmental aneuploidies to pinpoint “critical regions” on the two chromosomes, especially on chromosome 10^13–21^. More recent studies used genotyping arrays and high-throughput sequencing^22–25^. Position-based marker techniques were coupled with gene cloning techniques to identify possible culprit genes on chromosomes 10 and 7. One idea was to search for genes whose expression change correlates with the copy number change, and sometimes additional functional evidence was considered^7,21–24,26,27^. A related method was to seek genes that have frequent somatic mutations correlated with changes in expression^28^.

Analyses of chromosome 10 identified three regions that are most frequently lost, one on 10p and two on 10q^13,14,16,25,29–34^. In the region closest to the 10q telomere, the tumor suppressor gene *PTEN* was cloned^35–37^ and identified as a likely suspect in GBM pathogenesis^15,35,36^. The fact that *PTEN* is often the target of somatic mutations in the retained copy of chromosome 10 bolstered the evidence^38,39^, but a meta-analysis of over 10,000 patients suggested that *PTEN* loss is unlikely sufficient to explain all chromosome 10 losses^40^. Even though 10q is lost in approximately 80% of GBMs, the biallelic inactivation of *PTEN* occurs only in about 40% of cases, and GBMs without *PTEN* biallelic inactivation showed similar *PTEN* expression levels as samples in which 10q was not lost^41,42^. The gene *ANXA7* has been suggested as another possible culprit in a frequently deleted region of 10q closer to the centromere^43–45^. *ADARB2* and *KLF6* have been suggested as possible culprit genes in the critical region on 10p^25,46^.

The analysis of chromosome 7 gain has been less informative in terms of identifying culprit regions and genes because 7 gain usually covers the whole chromosome^7^, but some segmental CNAs of various intervals on chromosome 7 were detected^19^. The most commonly suggested culprit genes are *EGFR* on 7p^5,40,47,48^ and *MET* on 7q^48,49^. Several studies have claimed that specific gene pairs such as *PDGFA* on chromosome 7 and *PTEN* on chromosome 10^7^ or small sets of genes fully explain the co-occurrence of 7 gain and 10 loss^7,50–52^. The evidence relating the roles and relationships of *EGFR* and chromosome 7 gains is complex, however. EGFR is one of the most studied among the > 50 human receptor tyrosine kinases, but the mysteries of EGFR are still being elucidated^53^. *EGFR* amplifications often co-occurs with aberrant transcripts, especially a transcript denoted EGFRvIII, which lead to constitutive EGFR signaling^11^. *EGFR* has both kinase-dependent and kinase-independent pro-tumorigenic functions^53^. *EGFR* aberrations in GBM may manifest in at least six ways that are not mutually exclusive: i) activating point mutations, ii) genomic amplification, iii) chromosomal rearrangements, iv) autocrine signaling, especially between mutant forms of EGFR, such as EGFRvIII, v) intragenic duplication of the kinase domain, vi) *EGFR* fusions with pieces of other genes^53^. High-level *EGFR* amplifications may occur either on double minute chromosomes or via insertions of extra copies of *EGFR* on otherwise normal chromosome 7^54^. *EGFR* aberrations in GBM may be clonal or subclonal^54,55^. A study of 86 GBMs found that *EGFR* amplification occurs with higher probability in samples that have a gain of chromosome 7 (82.1%) compared to samples that do not (66.7%), but all four combinations for *EGFR* amplification or not and chromosome 7 gain or not were observed^56^, which qualitatively confirmed an earlier study done with older assays^57^. Chromosome 7 copy number gains to trisomy or moderate polysomy show no clear association with *EGFR* expression, although *EGFR* amplifications to double-digit numbers of gene copies do show a significant association with *EGFR* expression^58^. The mixed association results yield doubt that *EGFR* is the sole culprit gene among GBMs harboring a gain of 7p encompassing *EGFR*.

In summary, there is no small set of genes identified to date that can fully explain the notoriously frequent 10 loss/7 gain co-occurrence. Recent literature has supported the notion that aneuploidy patterns are shaped by the accumulation of multiple haploinsufficient and triplosensitive genes^59–61^. Therefore, in this work, instead of aiming to identify specific genes whose losses or gains on chromosome 10 and chromosome 7 can provide an explanation to the 10 loss/7 gain double event, we take a non-reductionist approach by analyzing chromosomes 10 and 7 as two large collections of genes whose protein products may interact, driving the 10 loss/7 gain combination. First using a rigorous mathematical model that we developed, we show that 7 gain after 10 loss is significantly more likely than the opposite case and that the less frequent order of 10 loss after 7 gain can be treated as random. Next, by analyzing patient tumor and cell line datasets, we provide an evolutionary explanation of why the combination of 10 loss and 7 gain aneuploidies is so prevalent in gliomas. Finally, by analyzing transcriptomic data from normal, non-cancerous human brain tissues, we provide a plausible reason why 10 loss and 7 gain co-occurrence happens preferentially in cancers originating in certain regions of the brain.

## RESULTS

### Analysis Overview

We use the phrasing “10 loss” or “7 gain” to denote loss of either arm (or the whole chromosome) of chromosome 10 or gain of either arm of chromosome 7, respectively (precise rules for calling an arm as lost or gained are provided in **Methods**). The terms *10 cnn* or *7 cnn* indicate a copy number neutral state where there is no arm loss and no arm gain of chromosome 10 or chromosome 7, respectively, compared to the modal ploidy (**Methods**).

Our analysis proceeds in four main steps:

1. *Mathematical Modeling:* We developed mathematical models to study the order of co-occurring aneuploidies. Applying these models, we showed that the probability of 7 gain occurring after 10 loss is significantly greater than 10 loss after 7 gain, finding that the less frequent sequence 10 loss after 7 gain can be treated as occurring by random chance.
2. *Data Mining for identifying Genetic Interactions:* We analyzed hundreds of genomic and transcriptomic samples from brain cancer patients and cell lines. Our aim was to identify a category of clinically significant genetic interactions that functionally compensate for the genes located on the lost arm (synthetic rescues). This step revealed that genes on chromosome 10, when lost, can be best functionally compensated by the upregulation of genes on chromosome 7, compared to genes on other arms. We also show that glioma patients with 10 loss and 7 gain events tend to have worse overall survival, testifying to enhanced underlying tumor fitness.
3. *CRISPR Screening for Fitness Effects:* Analyzing large-scale *in vitro* CRISPR essentiality screens in central nervous system (CNS) cancer cell lines, we show that the fitness effects of different possible sequences of chromosomal arm alterations further support the capacity of 7 gain to compensate for 10 loss.
4. *Normal tissue analysis:* Although some of our analyses in parts 1-3 are specific to brain cancer in the sense that they analyze brain cancer data, those analyses do not explain why the co-occurrence of 10 loss and 7 gain is so high in specific brain cancers (such as GBM) which preferentially arise from certain regions of the brain, but not in cancers from other body locations. Therefore, we analyzed expression data from thousands of normal non-cancerous samples from human brain tissues. In this analysis of non-cancer data, we provide evidence that the normal transcriptome state in the cortex or frontal cortex (which are common tissues of origin for GBM), are permissive of a 10 loss and 7 gain co-occurrence in cancers arising from these tissues.

### Chromosome 10 loss and chromosome 7 gain co-occurrence frequency and their associations with patient survival in gliomas

We began by analyzing over 39,000 patient tumors in the large-scale Progenetix database^62,63^, omitting tumors in the database that were also in The Cancer Genome Atlas (TCGA) cohort^64^, to quantify the relative prevalences of the 10 loss and 7 gain chromosomal events in glioblastoma in comparison to other loss-gain co-aneuploidy events occurring in other types of cancer. In this analysis, we considered p and q arms separately and focused on autosomes because Progenetix does not curate the sex of each subject.

We identified 1,280 significantly co-occurring chromosome arm loss-gain event pairs out of 31,122 possibilities (Fisher exact test, false discovery rate or FDR < 0.05; **Table S1**; **Methods**). Our findings are in line with previous work by Prasad et al.^65,66^, which examined the TCGA cohort^64^. We find a significant overlap between our identified co-occurring loss-gain events and theirs (Fisher exact test, Odds ratio (OR) = 15.0, P = 1.65e-39, **Methods**). Notably, we find that chromosomes 10 and 7 (either p or q arm) in GBM are the most significantly enriched co-occurring chromosome loss-gain events across all such co-aneuploidy events occurring in any cancer type (**Fig. 1A**; robustness studies for different parameters shown in **Fig. S5**). The numbers of GBM patient samples in the Progenetix data where loss or gain occurs on chromosomes 10 or 7 are summarized in **Fig. 1B**. Because we see a high enrichment of loss and gain CNAs in both arms (p/q) for both 10 and 7 chromosomes, in follow-up analyses we considered chromosome-arm and whole-chromosome events together, that is, an event is a 10 loss if either 10p or 10q arm (or both) is lost and an event is a 7 gain if either 7p or 7q (or both) is gained.

**Figure 1:**
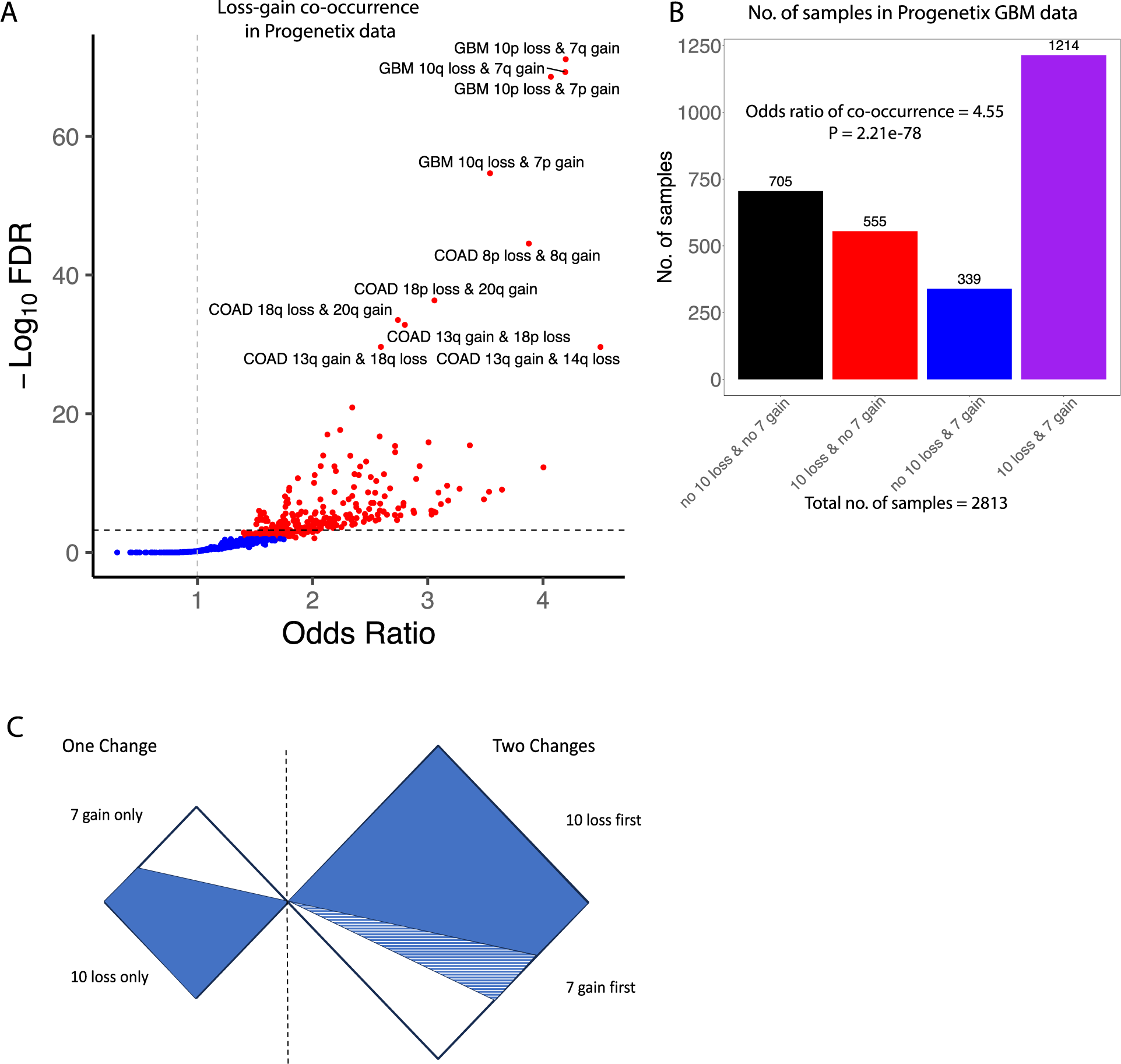
Chromosome arms co-occurring loss-gain event statistics and probabilistic analysis. **(A)** Plot charting the landscape of chromosomal arm loss-gain enrichments across various cancer types from patient tumors in Progenetix dataset. Each data point represents a chromosome arm loss/gain event for a specific cancer type (n = 5,886 data points in total), and the Odds Ratio (OR) and FDR values are obtained using Fishers exact test. The dashed vertical line is at OR = 1, and the dashed horizontal line is at FDR = 0.01. Events with FDR < 0.01 and OR > 1 are colored in red while the rest are in gray. In this figure, for the sake of clarity only events with at least 50 co-occurring cases in a specific cancer type are considered, and only a handful of top events are described (full results described in **Table S1**). There were no LGG cancers in the Progenetix data. **(B)** Bar plot showing the number of GBM patients in the Progenetix data with or without a 10 loss or 7 gain. 10 loss or 7 gain implies either the p/q arm is lost or gained respectively. The odds ratio and p-value for 10 loss and 7 gain co-occurrence using a two-sided Fisher exact test is shown. For the 705 cancers without 7 gain or 10 loss, an arm gain or loss was reported on some other chromosome indicating the tumor was aneuploid. Keywords – GBM: Glioblastoma multiforme, COAD: Colon adenocarcinoma. **(C)** Illustration of approximating the fraction of cooccurrences having 10 loss first by the number of tumors that have only 10 loss. The square to the left of the dotted line represents tumors with only one of 7 gain or 10 loss; the square to the right represents tumors with both. The solid blue area shows the proportion of each region for which 10 loss occurs first under the assumption that the events are independent. The blue striped area represents the greater number of tumors with 10 loss first followed by 7 gain that empirical evidence suggests would occur.

### Probabilistic model to estimate the probability of 10 loss occurring before or after 7 gain

Given the distribution of 10 losses and 7 gains that we found in the Progenetix data, we asked whether we could determine whether the loss of 10 loss or the gain of 7 were more likely to have occurred first. For the first part of the mathematical modeling, the intuition that a more frequent discrete genomic aberration (e.g., 10 loss could be a CNA, a cytogenetic breakpoint, a DNA somatic mutation, or other) occurs preferentially earlier but not exclusively earlier and this intuition has been formalized repeatedly in tumor phylogenetics^67–69^. Tumor phylogenetic studies often find the best partial order of aberrations via combinatorial optimization, but that approach does not prove that the best partial order is statistically significantly more likely that the second-best order for just two aneuploidies. The order of events matters for our subsequent analysis because we hypothesize that the aneuploidy that occurs second may compensate for the aneuploidy that occurs first. Intuitively, if there are many more tumors that have 10 loss without 7 gain than tumors that have 7 gain without 10 loss, this suggests that 10 loss is more likely to come first.

To study if 10 loss is likely to occur before 7 gain, one may compute the fraction of tumors with only a loss of 10 among all tumors for which one of these events occurs (either 10 loss or 7 gain but not both, **Fig. 1C**, see mathematical details in **Supplementary note 1**). Considering the counts of 7 gain and 10 loss in Progenetix, we find that fraction is 555/(555 + 339) ≈ 0.625 (based on the counts from **Fig. 1B**), which is different from the 1/2 expected (if the two aneuploidies occur in either order with equal probability) by a one-sided binomial test with a p-value of 2.50e-13. This indicates that 10 loss occurs significantly more frequently before 7 gain than the opposite order.

Having established that this is the preferred order, we turn to study why the co-occurrence of 10 loss and 7 gain happens more frequently than expected if the two aneuploidy events were happening independently without any functional compensatory forces in play. To this end, we develop a ‘mixture model’ in we which we posit two types of tumors that have 10 loss first and 7 gain second. One type has the events occurring at random and the other has the events occurring predictably in the order 10 loss first and 7 gain second. Intuitively, if these ‘predictably ordered’ tumors are frequent enough then they are sufficient in number to explain all the excess co-occurrence of 10 loss and 7 gain. This is merely a model, and we cannot distinguish which individual tumors behave randomly and which behave deterministically. The idea that a formal mixture of two models for tumor progression can explain the data better than a single model also been used repeatedly in tumor phylogenetics^70–73^. Our intent is to prove that a mixture of two models explains the data better than one model, but two models suffice to allow us to proceed to the next step of analysis, concerning synthetic rescue interactions.

Using the counts given in **Fig. 1B**, we compute that the excess probability is entirely attributable to 10 loss after 7 gain if the probability of 10 loss coming first is 0.625 (details of the computation are provided in **Supplementary note 2**; **Fig. S1**), which exactly matches the estimate given above (and detailed in **Supplementary note 1**) from the Progenetix observed data. Reassuringly, the data provided in Körber et al.^12^ suggest an empirical estimate of *x* ≥ 0.595, which is not significantly different from 0.625 (binomial test). The analysis of Körber et al. uses time-series data on tumors not in Progenetix, so there is no circularity or self-reference when we compare our data analysis based on Progenetix to the previous study. We conclude that all the excess probability of co-occurrence can be explained by tumors in which deterministically 10 loss is first and 7 gain is second. This additionally suggests that the opposite event of 7 gain first and 10 loss second can be treated as occurring by chance.

### Co-occurrence of 10 loss and 7 gain is associated with worse survival in glioma patients

We next analyzed clinical genomic data to ask whether 10 loss, 7 gain, and their co-occurrence, might be associated with the patients’ clinical outcome. We first analyzed the TCGA GBM and LGG patient data^41,74,75^ to study the effects of these events on patients’ survival. This analysis reveals that patients with 10 loss and 7 gain events have significantly shorter overall survival than patients whose tumors with 10 loss and 7 cnn, 10 cnn and 7 gain, 10 cnn and 7 cnn (**Figs. 2A**, **S2A-C);** our survival analysis differs substantially from the survival analyses in two previous publications^9,10^, which grouped the patients using other genomic features. Survival keeps worsening with each event in the following order: 10 cnn and 7 cnn (maximum survival), 10 cnn and 7 gain, 10 loss and 7 cnn, 10 loss and 7 gain (worst survival). These survival trends remain if we consider only patients without an *IDH1/2* mutation or GBM patients only (**Figs. S2D,E**). This testifies that 10 loss solely leads to a more aggressive form (lower survival) than 7 gain only, and 10 loss and 7 gain is the most aggressive state. This is in line with the notion that the co-occurrence of 10 loss and 7 gain increases the fitness value of tumors, in comparison to each of the individual aneuploidies.

**Figure 2:**
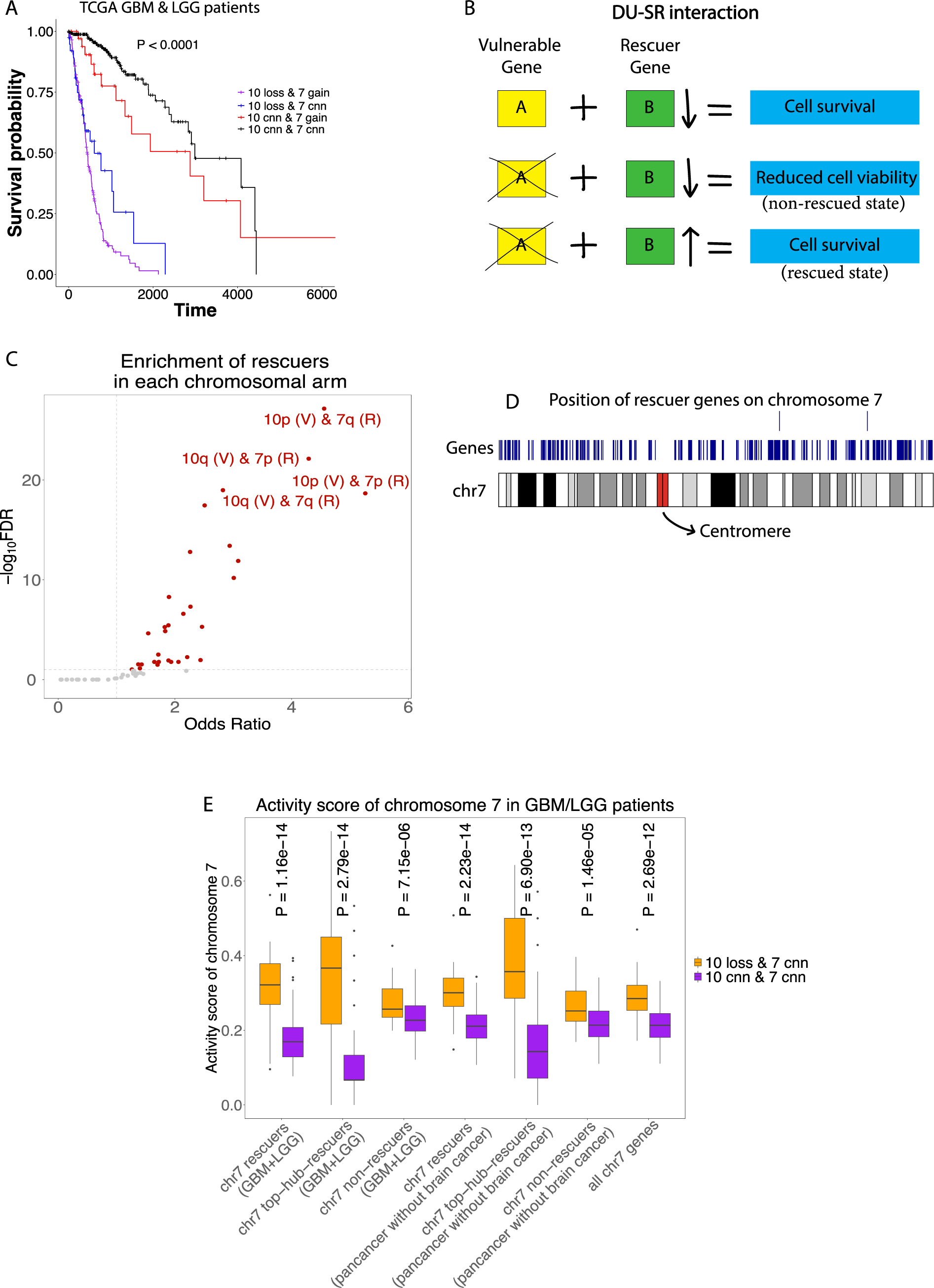
Evolutionary insights into the loss of chromosome 10 and gain of chromosome 7 in gliomas by analyzing patient tumor data. **(A)** Survival analysis (Kaplan-Meier curves) in TCGA GBM/LGG patients with the following occurrences: 10 cnn and 7 cnn (n=259), 10 loss and 7 cnn (n=38), 10 cnn and 7 gain (n=40), 10 loss and 7 gain (n=174). Log-rank test p-value (P) is shown. Pairwise survival analysis between patients with 10 loss and 7 gain with the three other groups are shown in **Figs. S2A-C**. **(B)** An illustration showcasing a DU-SR interaction between two genes A and B (adapted from Figure 1 in Sahu et al. 2019^79^). **(C)** Volcano plot showing the overlap enrichment of rescuer genes (from GBM-LGG DU-SR analysis) with the genes in each chromosome arm (Fisher exact test). The dashed horizontal line is FDR = 0.1, and the dashed vertical line is at odds ratio = 1. The rescuer genes are identified for all the (vulnerable) genes on a specific arm and the tested for overlap enrichment across all chromosome arms. Rescuers of vulnerable genes in 10p or 10q are more enriched in chromosomes 7p or 7q. **(D)** A plot of rescuer genes on 7 against ideograms of chromosome 7 generated using the karyoploteR package^82^. The full extent of each gene, from 5’ most exon in any transcript to 3’ most end of any transcript, is shown in blue; because of the scale of genes versus the scale of chromosomes these may appear as lines. On rare occasions, genes overlap, and when this occurs one of the genes is plotted in a second row above the ideogram. **(E)** Box plots showing activity scores on chromosome 7 between two groups of GBM/LGG patients: 10 loss and 7 cnn (n=38) versus 10 cnn and 7 cnn (n=259). Activity score is the fraction of a defined set of genes on chromosome 7 which have high expression (**Methods**). The defined set of genes are: (i) rescuer genes on chromosome 7 (chr7); (ii) top hub rescuers on chromosome 7 (i.e., those that map to ≥ 10 vulnerable genes on chromosome 10); (iii) genes on chromosome 7 which are not rescuers; (iv) all chromosome 7 genes. DU-SR network is computed using TCGA GBM+LGG data or TCGA pan-cancer data without using brain tumors. Keywords – GBM: Glioblastoma multiforme, LGG: brain lower grade glioma; chr7: chromosome 7; 10 cnn or 7 cnn: copy number neutral state for chromosomes 10 and 7, respectively; ‘&’ implies ‘and’.

### Genes on chromosome 10 are enriched for functional rescuer genes on chromosome 7, testifying for 7 gain capacity to compensate for 10 loss

Loss of many genes on a chromosomal arm is likely to be detrimental to cancerous cells^60,76–78^. We set out to study the hypothesis that at least some negative effects for gliomas due to the loss of many genes on chromosome 10 could be rescued by gains of genes on chromosome 7. We address this by identifying clinically relevant genetic interactions known as synthetic rescues.

*Synthetic rescue (DU-SR) interactions* refer to a form of functional interplay in which the detrimental impact on cell fitness caused by the inactivation of a specific gene (referred to as the *‘vulnerable gene’*) is compensated by upregulation of another gene known as the *‘rescuer gene*’^79^ (**Fig. 2B**). We previously published and validated a computational pipeline called ‘IdeNtification of ClinIcal Synthetic Rescues in cancer’ (INCISOR), which mines genomic, transcriptomic, and phenotypic data (such as patient survival) from hundreds of cancer patients and cell lines to identify DU-SR interactions that may be clinically relevant in patients^79,80^ (**Methods**). INCISOR first identifies DU-SR interactions by mining hundreds of *in vitro* essentiality screens from cancer cell lines, and then filters those interactions by looking for positive selection and worsening survival association in patients’ tumors, and for interactions between gene pairs with high phylogenetic similarity across many divergent eukaryotic species^79^. Many of the predicted clinically relevant SR interactions were validated through experimental *in vitro* screens and by their ability to predict cancer drug response in patients^79,81^. While the terms ‘vulnerable’ and ‘rescuer’ are used for historical reasons (since usually rescuer genes are identified for ‘vulnerable’ drug targets), INCISOR identifies any gene pairs where the downregulation of a gene and the upregulation of the partner gene is beneficial for cancer, over and above their individual gene effect^79^.

To investigate the role of DU-SR interactions in gliomas, we applied INCISOR to analyze 664 TCGA GBM and LGG patients to predict genome-wide clinically relevant DU-SR interactions for the (vulnerable) genes on chromosome 10 (**Methods**, **Table S2**). Remarkably, we found that chromosome arms 7p and 7q have the highest enrichment of rescuer genes for the vulnerable genes in 10p and 10q among all chromosome arms, in comparison to what is expected by chance (overlap enrichment test using Fisher exact test, FDR < 10^-19^, **Fig. 2C**, **Table S3A**, **Methods**). We also observe that the rescuers on chromosome 7 are positionally spread throughout the chromosome (**Fig. 2D**), which may explain why gains of chromosome 7 in brain cancer often involve whole chromosome arms rather than smaller regions. This analysis testifies that the loss of genes on chromosome 10 in gliomas could be compensated by the amplification of a high number of rescuer genes on chromosome 7.

To avoid circularity concerns that the DU-SR results were only due to the high prevalence of brain cancers with 7 gain and 10 loss in the data, we repeated the DU-SR analysis on TCGA GBM and LGG patients by explicitly removing an INCISOR step that looks for positive selection of gene pairs with high expression or copy-number variation. Reassuringly, this analysis yielded similar results (**Fig. S3**; there are no circularity concerns in any of the other steps of INCISOR). Second, we performed the DU-SR prediction analysis on the pan-cancer TCGA data *after removing GBM and LGG tumor samples* (**Methods**, **Table S4**). Remarkably, the strong enrichment of rescuer genes in chromosome 7 for the vulnerable genes in chromosome 10 persisted (**Table S3B**).

Next, we compared gene expression patterns between tumor samples from TCGA GBM/LGG patients who have lost chromosome 10 with those with 10 cnn (chromosome 7 is in a copy number neutral state in both groups). We asked whether genes on chromosome 7 would be over-expressed when chromosome 10 is lost. We formulated an “activity score” based on the proportion of a defined set of genes on chromosome 7 that have high expression levels when chromosome 10 is lost (**Methods**), uncovering an overall increased expression of genes on 7 (i.e., higher activity scores) when chromosome 10 is lost. Importantly, this heightened expression is pronounced amongst the predicted rescuer genes— especially those capable of rescuing a multitude of vulnerable genes on chromosome 10—compared to non-rescuer genes (**Fig. 2E**). In other words, when chromosome 10 is lost, even when chromosome 7 is not gained, *the rescuer genes that lie on 7 are highly expressed*, testifying to the possible role of rescuer genes on chromosome 7. Repeating this analysis while excluding brain tumors reassuringly yielded congruent results (**Fig. 2E**).

Our analysis overall predicts 237 vulnerable genes and 272 rescuer genes on chromosomes 10 and 7, respectively. Reassuringly, some of the previously suggested candidate genes *PTEN* and *ADARB2* were among the predicted vulnerable genes on chromosome 10 (see **Introduction**), whereas some of the genes previously reported to drive 7 gain, such as *EGFR, MET, BRAF,* where among the identified rescuers (**Table S2**). A ranked list of rescuers (based on the number of vulnerable genes they interact with) is provided in **Table S2D**, along with a pairwise pathway enrichment analysis (**Supplementary note 3, Fig. S4, Table S5**). Given prior suggestions that the loss or gain of these chromosomes could be driven by a few key genes, we conducted the following sensitivity analysis. After removing previously reported well-known genes among those present in DU-SR network (*PTEN, ADARB2* from chromosome 10 and *MET, BRAF, EGFR* on chromosome 7), we still observed that the rescuer interactions for the vulnerable genes on chromosome 10 are highly enriched with genes in chromosome 7 (FDR < 10^-19^, **Table S3C**). This suggests that the chromosomal events of 10 loss and 7 gain are likely orchestrated by a broader network involving many genes, beyond just a few previously reported genes.

### Essentiality screens in CNS cell lines further testify to the fitness benefits of 7 gain in the presence of 10 loss

Independent from our synthetic rescuer based analysis, we next turned to study the potential selective advantage of 10 loss and 7 gain by analyzing the DepMap dataset of gene essentiality screens (via CRISPR-based knockouts) in IDH-wildtype CNS cancer cell lines^83,84^. First, we focused on CNS cell lines with 10 loss and partitioned them into 7 gain and 7 cnn groups, and then compared the relative essentiality between the two groups for the genes on chromosome 10 and 7. We find that overall, chromosome 10 genes are likely to be less essential in the 7 gain group than in the 7 cnn group which is in line with our findings in the brain tumor analysis (two-sided Fisher exact test, P = 0.0034 in comparison to the all-gene analysis; **Methods**; **Fig. 3A, Table S6A**); however, chromosome 7 genes tend to be more essential in the 7 gain group (P = 0.0019, **Fig. 3A**). Besides, chromosome 7 genes also tend to be more essential in 10 loss and 7 gain cell lines than in the 10 cnn and 7 gain cell lines (P = 1.11e-23, **Fig. 3B, Table S6B**). In contrast, when comparing 7 gain vs 7 cnn groups in 10 cnn cell lines, chromosome 7 genes tend to be less essential with 7 gain (two-sided Fisher exact test, P = 9.75e-18 in comparison to the all-gene analysis; **Fig. 3C, Table S6C**). Since 7 gain in the presence of 10 loss is associated with decreased essentiality to chromosome 10 genes and increased essentiality to chromosome 7 genes, this suggests an evolutionary pressure to maintain 7 gain in the presence of 10 loss, consistent with an associated selective advantage. Furthermore, we analyzed a CRISPR screen of recently developed isogenic system of RPE1 cells molecularly engineered to have distinct single whole-chromosome trisomies^85^. The chromosome 10 genes that were essential in the near-diploid clone (RPE1-SS48) were significantly less essential in the clone with trisomy 7 (RPE1-SS6), but not in the clone with trisomy 8 (RPE1-SS119) (**Fig. 3D; Methods**). The RPE1 cell line analysis further supports the likely evolutionary benefit of gaining chromosome 7 after the loss of 10.

**Figure 3:**
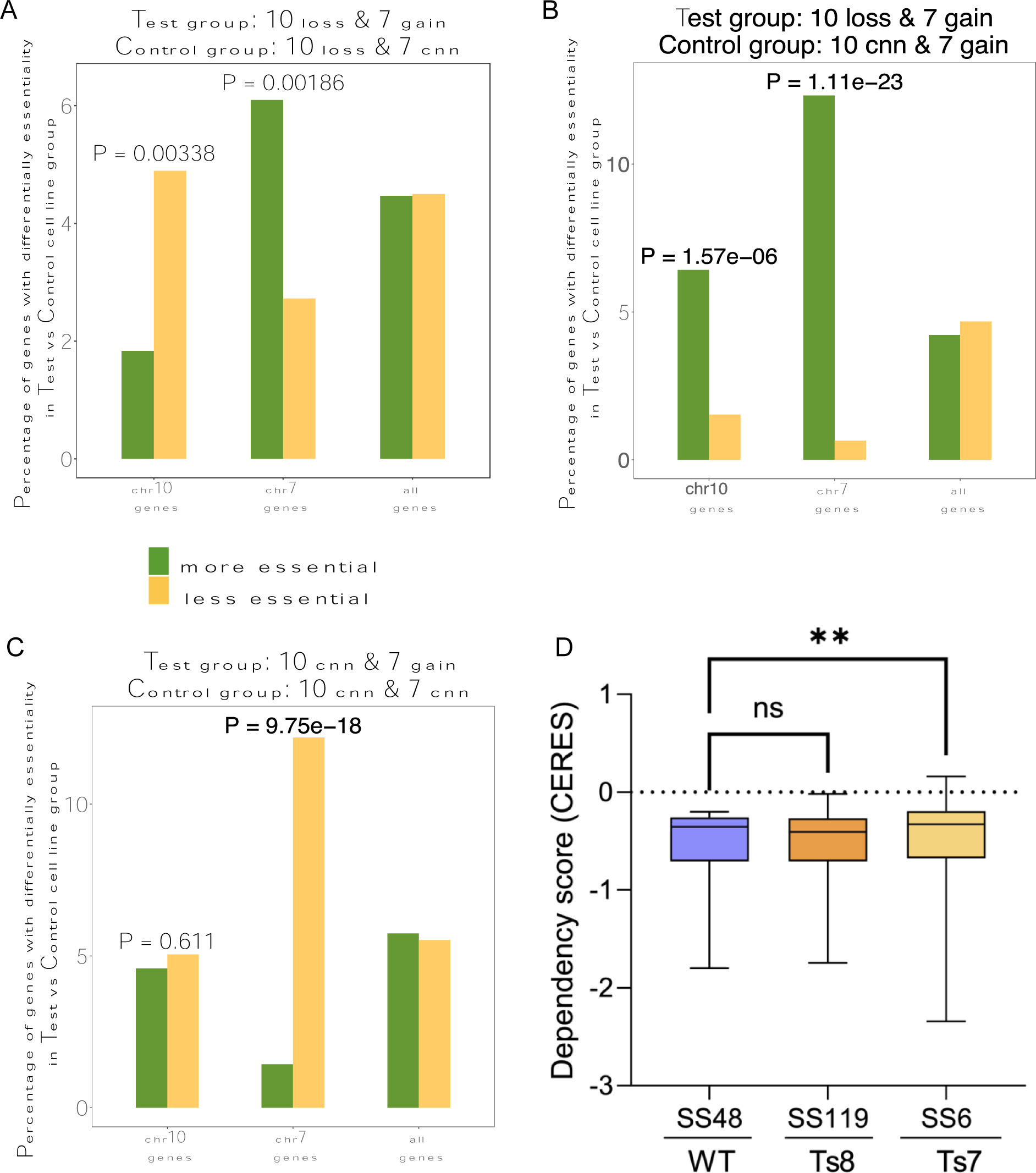
In vitro essentiality analysis associated with 10 loss and 7 gain on central nervous system (CNS) cancer cell lines. **(A-C)** Bar plots showing the percentage of more (green) or less (yellow) differentially essential genes between two groups of CNS cell lines: **(A)** test group (10 loss and 7 gain, n=49) vs control group (10 loss and 7 cnn, n=8) of CNS cell lines; **(B)** test group (10 cnn and 7 gain, n=22) vs control group (10 cnn and 7 cnn, n=8); **(C)** test group (10 loss and 7 gain, n=49) vs control group (10 cnn and 7 gain, n=22). **(D)** Box plots comparing the dependency scores of chromosome 10 genes between a near-diploid RPE1 clone (SS48), a clone with trisomy of chromosome 7 (SS6) and a clone with trisomy of chromosome 8 (SS119). Only genes that are defined as essential in SS48 (CERES<-0.2) are considered for this analysis.

### Why do 10 loss and 7 gain happen in certain regions of the brain rather than other regions?

Among the many different kinds of brain cancers, 10 loss and 7 gain mainly co-occur in GBMs, which are more likely to arise in the supratentorial regions of the cerebral hemisphere (e.g., cortex or frontal cortex)^86,87^ than in other regions of the brain. While previous sections of this paper focused on studying cancer samples, here, to understand why this specific co-occurrence is likely to happen in cancers arising from certain regions of the brain, we analyzed gene expression data from 2,642 samples from 13 types of normal non-cancerous brain tissues from the Genotype-Tissue (GTEx) dataset^88^. The GTEx analysis is motivated by a previous study that showed that the inferred synthetic lethality activity in normal tissues can explain tumor suppressors’ role in cancers arising more frequently in specific tissues vs other tissues^89^.

To recap the parts of the preceding analyses relevant to this subsection, we have applied INCISOR to identify genetic interactions (DU-SR network) in chromosomes 10 and 7 where the inactivation of gene on chromosome 10 and its activation of its partner gene on chromosome 7 is beneficial for cancer. To study the potential effects of the INCISOR-derived DU-SR network (derived from brain tumor data) in normal non-cancerous tissues in analogous fashion to the analysis conducted in (Cheng, Nair et al., 2021)^89^, we defined a measure called *“cancer synthetic rescue (cSR) load*”, which is a quantitative measurement of naturally occurring ‘potent’ DU-SR interactions in normal samples between gene pairs on chromosomes 10 and 7 (**Methods**; an overview of the approach using a mock example is shown in **Fig. 4A**). cSR load in a single sample is defined as the fraction of DU-SR pairs on chromosomes 10 and 7 in which the gene on chromosome 10 has low expression (i.e, inactive) and the partner gene on chromosome 7 is not lowly expressed and hence likely to be active. Tissue cSR load (across all samples from that tissue) is the median value of all single-sample cSR loads in a specific normal tissue.

**Figure 4:**
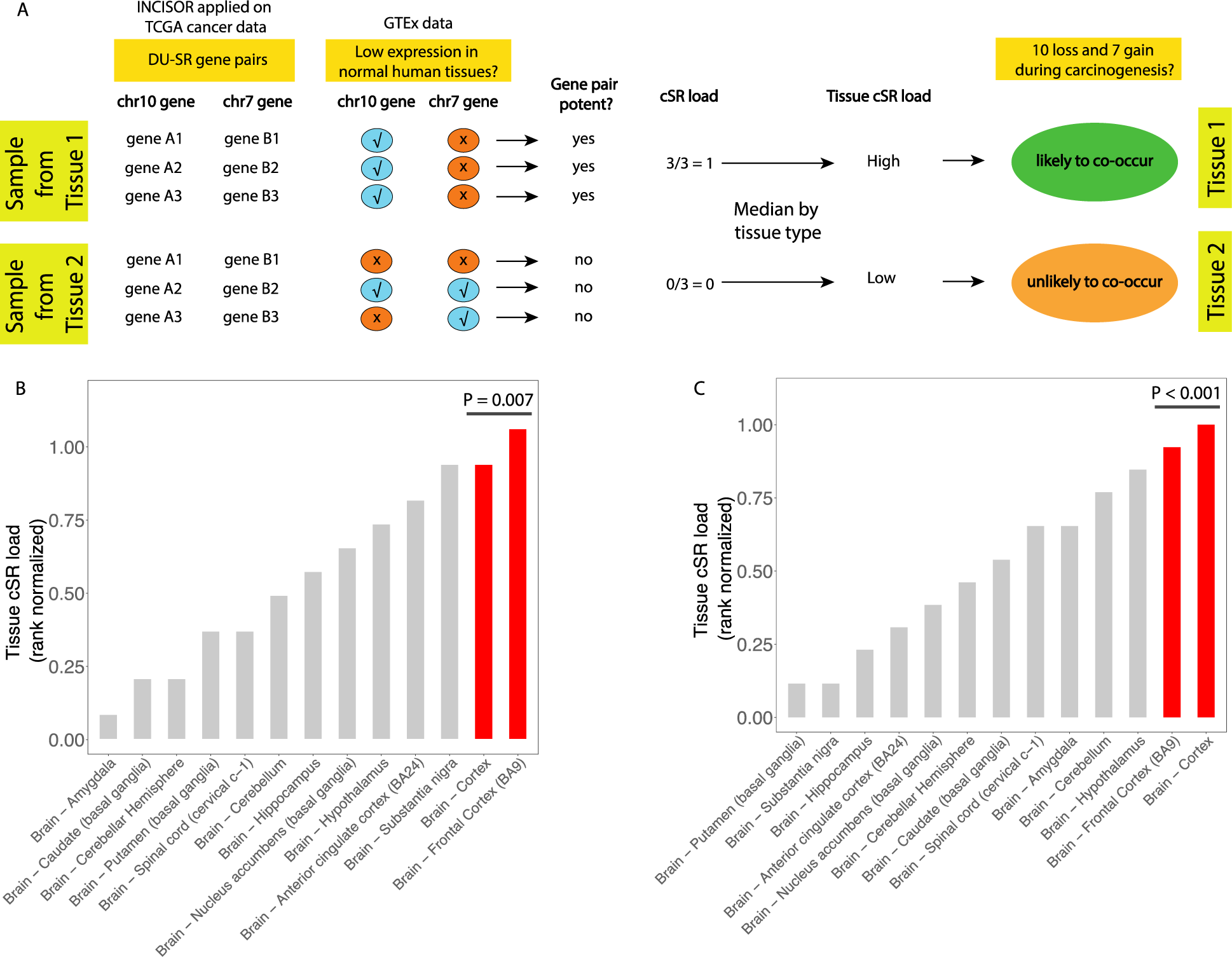
Normal non-cancerous brain tissue analysis. **(A)** Toy example demonstrating the computation of cancer synthetic rescue (cSR) load. chr10 and chr7 stands for chromosomes 10 and 7 respectively. **(B, C)** Bar plots showing the variation of tissue cSR loads (rank normalized across tissues) across various brain tissues in the GTEx database, using DU-SR networks derived from: **(B)** brain cancer data; **(C)** pan-cancer data while excluding brain tumors. Frontal cortex and cortex tissues are shown in red as they are most relevant to GBMs. Empirical p-values (P) of both frontal cortex and cortex being ranked as the top two based on their tissue cSR loads in comparison to random controls are shown (**Methods**).

Since 10 loss and 7 gain are early events in cancer development, we hypothesized that normal tissues with higher tissue cSR loads are more likely to develop 10 loss and 7 gain co-occurrence during the process of carcinogenesis, since the latter testifies to the compensatory potential of cell survival after the loss of 10. As in ^89^, the DU-SR interactions on chromosomes 10 and 7 are derived from cancer and not normal samples, but we hypothesize that they could become functionally relevant as cancer develops.

Following this line of thought, we computed the Tissue cSR loads of 13 normal non-cancerous brain regions in GTEx samples. Our results show that among all these 13 brain tissues, the frontal cortex and cortex tissues have the highest tissue cSR loads, and notably, these are the two most relevant tissues to GBM in the GTEx cohort (**Fig. 4B**). Control experiments using randomly generated cSR networks did not show a similar pattern (empirical p-value of both these two tissues being ranked in top two based on their cSR load in comparison to random controls is 0.007; **Fig. 4B; Methods**). We repeated this analysis by computing DU-SR interactions (on chromosomes 10 and 7) in a pan-cancer manner by explicitly removing any brain tumor data. Reassuringly, we again see the cortex and frontal cortex as the two tissues with the two highest cSR loads (empirical P < 0.001; **Fig. 4C; Methods**). These results suggest that the pre-existing transcriptomic state in the cortex and frontal cortex might allow for tolerating and compensating for the loss of 10 by rescuer genes that are active on 7, whose active upregulated transcriptional state is then further fixated by the gain of 7, which they drive and select for.

## DISCUSSION

The mystery behind the co-occurring loss of chromosome 10 and gain of chromosome 7 in many gliomas has been investigated since the late 1980’s. Prior studies have tried with limited success to explain these frequently co-occurring events focusing on a few driver genes as culprits. In contrast, our comprehensive genome-wide analysis suggested a different perspective. Acknowledging that it is still likely that a few drivers play important roles in 10 loss and 7 gain co-occurring gliomas, our analysis suggests that this phenomenon is further orchestrated by a complex interaction of many genes residing within these two chromosomes, where increased expression of multiple rescuer genes on the gained chromosome 7 compensates for the down-regulation of multiple vulnerable genes on the lost chromosome 10.

Analyzing data from Progenetix, we developed a probabilistic model that elucidated the probability of these chromosome co-occurring events, by estimating the probability of 7 gain after 10 loss and vice-versa. In particular, we found that 7 gain after 10 loss is the more likely event, and that the opposite event can be treated as occurring by random chance. Next, we found that while it remains possible that the loss of chromosome 10 in many GBM/LGG tumors is driven by a few drivers, this loss may be enabled by the tumor’s ability to readily compensate and mitigate the adverse effects it triggers. Even when a few driver genes underlie the recurrence of a lost chromosome it is widely expected that the loss of many other genes on the same chromosomal-arm would be detrimental, thereby reducing the positive selection towards that chromosome-arm loss ^60,76–78^. We show here that gaining chromosome 7 after the loss of 10 increases the fitness advantage conferred to GBM/LGG tumors by 10 loss, primarily through intricate genetic rescue interplays between genes on these chromosomes. Even during instances where a specific gene loss on chromosome 10 is neutral or beneficial for cancer (instead of being detrimental), INCISOR can identify partner genes on chromosome 7 whose activation can provide a synergistic benefit to cancer cells. Further independent investigation in IDH wild-type CNS cancer cell line essentiality screens reinforced the conclusion the gain of 7 in the presence of 10 loss is associated with a selective advantage. Lastly, we analyzed the transcriptomic data from normal non-cancerous human brain tissues and found that the preexisting transcriptomic state of the cortex and frontal cortex make them predisposed to cancers with 10 loss and 7 gain co-occurrence due to the high activity of rescuer genes on chromosome 7.

Of course, there are also gliomas where 10 loss alone happens without the gain of chromosome 7. Interestingly, we do see that in such cases, the rescuer genes we identified on chromosome 7 tend to have higher expression than tumors where 10 is not lost, thereby testifying to the compensatory role of the genes on chromosome 7.

Our findings address another puzzle about the co-occurrence of 10 loss and 7 gain. Previous studies noted that the losses on chromosome 10 are often segmental, but the gains on chromosome 7 more often encompass the entire chromosome, albeit with many exceptions^7,10,16,48^. One explanation suggested previously for this distinction is that an epistatic oncogenic benefit is achieved when both the oncogenes *EGFR* and *MET* and other genes on chromosome 7 (e.g., *HGF* and *PDGFA*) are gained^90^. Since these two gene pairs located far apart and near the two chromosome 7 telomeres, the evolving cancer is likely to gain the entire chromosome 7. Extending upon this earlier suggestion, we find that the predicted rescuer genes on chromosome 7 are distributed throughout the entire chromosome, pointing to a potential compensatory role of the gain of the whole chromosome.

Beyond providing a comprehensive evolutionary account of the frequent 10 loss/7 gain co-occurrence, our analysis provides specific predictions of genes involved. Of special interest are major rescuer genes that reside on chromosome 7, whose targeting may have a therapeutic potential, if further corroborated in future gene-specific experimental studies. The experimental testing of such predictions is a tall order on its own and is out of the scope of the current investigation, which is focused on presenting a first of its kind holistic evolutionary explanation of this fundamental co-occurring event. However, to facilitate such a future exploration, we cataloged a prioritized list of vulnerable/rescuer genes that reside on chromosome 10/7 (**Tables S2, S4**).

In conclusion, this analysis presents a new multi-pronged approach to analyze co-occurring aneuploidy events in cancer, shedding new light on a long-contemplated chromosomal co-occurrence event in gliomas. Notably, it could be applied in future studies of other common co-occurring aneuploidies across many cancer types.

## METHODS

### Analysis of Progenetix data

We downloaded the metadata from tumors from 118,238 patients from the Progenetix^63^ database on January 5, 2023 (https://progenetix.org/). The metadata included histological diagnosis identifiers, which are specified using a hierarchical system of National Cancer Institute (NCI) Thesaurus Terms. We searched this ontology of terms to identify patients that could be unambiguously assigned to 24 TCGA cancer types. Specifically, we used the “rols” package in GNU R to download and search the ontology. NCI Terms and TCGA cancer types are not exactly one-to-one matches, but GBM was an unambiguous diagnosis in Progenetix.

We then downloaded segmental copy-number data from Progenetix, also on January 5. A total of 50,392 patients could be unambiguously identified with a unique TCGA type and had CNV data. We further eliminated those samples in the TCGA projects and randomly selected one tumor sample from each person having more than one sample, which left 39,085 persons and 39,085 samples. We used only tumor samples, not matched normal, but some persons had more than one tumor sample. We chose one primary tumor sample per person randomly, yielding 39,085 tumors. Samples in Progenetix sometimes have more than one set of CNV calls, possibly due to multiple sequencing runs, but this only affects individuals in TCGA. The segment data in the Progenetix database do not distinguish ambiguous regions from calls of neutral copy number. Nor does the technology used for much of the Progenetix database, comparative genomic hybridization (CGH), identify the ploidy of the sample. It records deviations from neutral copy-number for that sample.

### Calling arm changes from Progenetix data

We identified arm changes using the following rules. We ordered the segments between the telomere of the arm and the centromere by position. Segments do not overlap, so there is no ambiguity in this order. We treated any segments not called by Progenetix for which

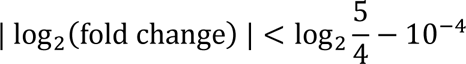

as having neither a gain nor loss call. The constant 5/4 was selected because it represents a gain on a ploidy background of 4 (the actual ploidy is unknown). The tolerance 10^−4^ compensates for the limited precision with which Progenetix saves its fold change numbers.

The most telomeric number change in this filtered ordered list determined the putative direction of arm change and must have been called within the telomeric 5 Mbp region of the chromosome arm. We then scanned the list from telomeric end to centromere, counting any contiguous sequence of segments either called in the opposite direction or having no call as a single gap. If the copy number change could be extended toward the centromere allowing at most 3 gaps and the number of bases called in the extension (not the length of the extension) covers 60% of the chromosome arm, then that arm was considered to be lost or gained. Progenetix does not distinguish between calls of copy number neutrality and regions that could not be called, which makes it impossiible to use a threshold derived from the percentage of bases in the chromosome. We found a threshold that 60% of an arm’s bases be gained or lost to be a reasonable value because it captured large losses on the telomeric end of an arm while allowing for large uncalled regions. We experiment with a stricter 80% threshold (which was the threshold used by from Taylor et al.^91^ for the TCGA dataset), but it did not result in a subjective difference in GBM (**Fig. S5A,B**). Using a rule that gain or loss of a chromosome arm was recorded a gain or loss of a chromosome, we performed a similar series of Fisher exact tests and once again found that loss of 10 with gain of 7 stood out as highly significant and the most strongly significant event (**Fig. S5C**).

### Calling chromosome loss/gain co-occurrences

To test for all co-occurring chromosome arm changes, for each cancer type, we used a Fisher exact test to determine whether the pair of arm changes occur more often than by chance given the rate of occurrence of the individual arm changes. Tests were one-sided because we only kept cooccurrences that happen often than expected. We only included samples for which there was evidence of aneuploidy, which we define here as the evidence of an arm change on any (non-acrocentric) autosome. We only consider loss/gain pairs because the CGH technology is relative to overall chromosome content, and we can be most confident that a loss/gain pair, having a change in each direction, reflects true events and not changes in the content of unrelated chromosomes. We use the FDR correction to compensate for multiple hypothesis testing^92^.

### Comparison of Progenetix analysis with previously published data

We compared our results to loss/gain pairs in Supplementary Table 4 in Prasad et al.^65,66^. We consider only loss/gain pairs, and therefore take the cancer specific pairs and p-values from Prasad et al, restrict them to the 22 TCGA cancer types we were able to identify, and recompute the FDR correction based on the pairs retained. From our own list, and the list of Prasad et al., we keep only those items with FDR-adjusted P ≤ 0.05. To test whether the list in Prasad et al. is enriched within our own, we use a single two-sided Fisher exact test for which we consider the universe to be all possible autosome arm gain/loss pairs among the 22 cancer types, ignoring the short ends of the five acrocentric chromosomes (13, 14, 15, 21, 22).

### DU-SR analysis

INCISOR is a computational method^79^ that identifies clinically relevant SR interactions that are supported by all these four steps to predict DU SR gene pairs.

a. Essentiality Screens: INCISOR mines essentiality data from hundreds of pan-cancer cell lines^84,93–97^ to identify genes that, when knocked down, can be rescued by the upregulation of another gene. In each gene pair, the first gene is called the vulnerable (V) gene and the second gene is called the rescuer (R) gene.
b. Survival of the Fittest: INCISOR uses hundreds or thousands TCGA patient tumor (genomic and transcriptomic) data to find SR pairs that appear more frequently in their rescued state (i.e., gene V is inactive and gene R is specifically upregulated), indicating positive selection.
c. Patient Survival: Using a stratified Cox proportional hazard model, INCISOR selects SR pairs (when gene V is inactive and gene R is upregulated) linked to worse patient survival, accounting for various confounding factors.
d. Phylogenetic Screening: Finally, INCISOR picks SR pairs with high phylogenetic similarity as the most likely candidates.

More details regarding the INCISOR pipeline can be found in Sahu et al.^79^. We ran INCISOR on 664 TCGA GBM and LGG patient samples and on hundreds of pan-cancer cell lines at FDR < 0.1 to identify DU-SR interactions in brain tumors. We also ran INCISOR on 8,085 TCGA pan-cancer patient tumor samples. For the pan-cancer analysis, we removed brain cancers from both cell line and patient datasets to avoid identifying brain-specific synthetic rescues which should be enriched in the previous TCGA test. Given the larger sample sizes a more stringent threshold of FDR-adjusted P < 0.05 was used in the pan-cancer analysis to identify DU-SR interactions.

### Enrichment analysis of rescuers on a specific chromosome

For all the genes on specific chromosome arms (e.g., chromosome arm 10q), we predicted DU-SR interactions in a genome-wide manner. Then, we looked for enrichment of rescuer genes among the genes on some chromosome (e.g., chromosome 7) and checked if they occurred more often on that chromosome than expected by random chance (Fisher exact test). We did a similar enrichment analysis for each chromosome arm. Statistical tests were corrected for multiple hypothesis testing using the FDR (false discovery rate) correction^92^.

### Cell line essentiality analysis

We used CRISPR gene-knockdown essentiality screen data from DepMap^84^ for our cell line essentiality analysis. Arm change event information for cell lines was taken from previous literature^85,98^. We then divided CNS cancer cell lines into two groups based on arm-level copy-number events. All CNS cell lines in the essentiality screen were wildtype for *IDH1/2* mutations. Genes on either chromosome 10 or chromosome 7 with differential essentiality between the two groups are identified using a one-sided Wilcoxon rank-sum test (P < 0.05). A similar analysis is also done by considering all genes. The number of more/less essential genes obtained for a specific chromosome or arm is then compared with the corresponding numbers obtained via the all-gene analysis using a two-sided Fisher exact test.

For the analysis of the isogenic RPE1 clones, CERES dependency scores of 3 clones – RPE1-SS48 (diploid), RPE1-SS6 (trisomy 7) and RPE1-SS119 (trisomy 8) – were retrieved from Zerbib et al.^85^. Genes with CERES score < −0.2 in the WT clone SS48 were included in the analysis. A non-parametric paired ANOVA was performed to determine the difference between the clones.

### Activity scores

Arm level copy-number data for TCGA patient tumors were sourced from Taylor et al.^91^. Taylor et al. considered chromosome arms with ≥ 80% change as aneuploid (value of +1 for gain or −1 for loss) and <20% as non-aneuploid (value 0) (more details in their paper^91^). Gene expression levels were categorized as high or low based on percentiles across all TCGA GBM+LGG patients: > 67th percentile for high expression and ≤ 33rd percentile for low expression. An ‘Activity Score’ was computed as the fraction of genes on chromosome 7 with high expression within specific gene subsets: (i) rescuer genes, (ii) top hub rescuer genes (those that mapped to ≥10 or ≥70 vulnerable genes on chromosome 10 using TCGA GBM+LGG or pan-cancer data, respectively), (iii) non-rescuer genes on chromosome 7, and (iv) all genes on chromosome 7.

### Normal non-cancerous brain tissue analysis

We used RNA-seq TPM data (log-scale) from 13 GTEx brain tissues containing 2642 normal non-cancerous samples. For each tissue, we further rank normalized (from 0 to 1) the expression data first across genes for every sample, and then across samples for each gene. A gene was considered to have low expression in a sample if its expression was less than or equal to the 33rd percentile of its expression across all samples in that tissue. The DU-SR network was derived using either TCGA brain cancer (GBM, LGG) data or using pan-cancer data without brain tumors. cSR load in a normal sample is defined as the fraction of DU-SR pairs (out of all DU SR pairs on chromosomes 10 and 7) where the vulnerable gene on chromosome 10 has low expression (inactive) and the rescuer gene on chromosome 7 does not have low expression (active). Tissue cSR load was computed as the median value of all sample specific cSR loads in a specific normal tissue. These tissue cSR loads were computed for all 13 brain tissues and rank normalized (from 0 to 1).

Control experiments were carried out by first randomly generating 1000 ‘cSR’ (sub)networks of the same size (i.e., same number of gene pairs) as our original network from a large network of all possible distinct gene pairs from across the genome. As before, rank normalized random-cSR loads were computed for all 13 brain tissues for each random network. An empirical p-value of both these two tissues having the two high values they have in the observed data based on their cSR load in comparison to random controls was computed as follows: the number of random networks in which the rank normalized random-cSR loads of both these two tissues (frontal cortex and cortex) are greater than or equal to the minimum observed rank normalized cSR loads of either one of these two tissues (using the actual cSR network) divided by 1000.

## DATA/CODE AVAILABILITY

The R code used to generate the main results in this paper has been made available for reproducibility: https://hpc.nih.gov/~Lab_ruppin/code_for_paper_braincancerproject.zip

## AUTHOR CONTRIBUTIONS

N.U.N. and E.R. conceived the research study. N.U.N., A.A.S., E.M.G. did most of the analysis with help from J.Z., G.L. The mathematical models were developed by A.A.S. and E.M.G. with feedback from N.U.N., E.R., K.C. Valuable insights into the methodology was provided by K.C, A.D.S., E.D.S., K.D.A., U.B-D. The results were interpreted by N.U.N., A.A.S., E.M.G., K.C., K.D.A., U.B-D, E.R. N.U.N., A.A.S., E.M.G., U.B-D., E.R. wrote the paper with inputs from all authors. All authors have read and approved the manuscript.

## ACKNOWLEDGEMENTS

This research was supported in part by the Intramural Research Program of the National Institutes of Health (NIH), National Cancer Institute, and the Center for Cancer Research. Work in the Ben-David lab was supported by the was supported by the European Research Council Starting Grant (grant #945674 to U.B-D.). This work used the computational resources of the NIH HPC Biowulf cluster (http://hpc.nih.gov). We thank Dr. Mark R. Gilbert, Dr. Orieta Celiku, Dr. Kun Wang, Mr. Neel Sanghvi for their feedback and help on this work. The results shown here are in part based upon data generated by the TCGA Research Network: https://www.cancer.gov/tcga.

## Declaration of generative AI and AI-assisted technologies in the writing process

During the preparation of this work the authors used ChatGPT 4 in order to improve the quality of writing for some of the sentences in this paper. After using this tool, the authors reviewed and edited the content as needed and takes full responsibility for the content of the publication.

