## Supplementary notes and supplementary figures for "Chromosome 7 to the rescue: overcoming chromosome 10 loss in gliomas"

### Supplementary material for paper "Chromosome 7 to the rescue: overcoming chromosome 10 loss in gliomas"

**SUPPLEMENTARY NOTES**

**1. More tumors exhibit loss of 10 before gain of 7 than the reverse.**

We use the terms '10 loss' or 'del10' interchangeably to denote loss of chromosome 10 (either p or q arm'); and '7 gain' or 'gain' interchangeable to denote gain of chromosome 7 (either p or q arm'), respectively. When we write about a gain of 7 before a gain of 10, or a loss of 10 before a gain of 7, we are referring to two time points: the point at which the first copy number change occurred and the point at which the tumor sample was collected. Let us call these time points $i-1$ and $i$ respectively. We introduce the notation $\mathrm{gain7_{i-1}}$ and $\mathrm{gain7}_i$ indicate that the copy number of 7 is higher than the modal ploidy at times $i-1$ and $i$, respectively. Let $\mathrm{del10}_{i-1}$ and $\mathrm{del10}_i$ similarly indicate a copy number of 10 less than the modal ploidy. We assume $\mathrm{gain7_{i-1}}$ implies $\mathrm{gain7}_i$; reversal of copy number changes is not permitted in this model. Similarly, $\mathrm{del10}_{i-1}$ implies $\mathrm{del10}_i$. In our model it also holds that exactly one of the two events happened first, no matter how small the time between the events.

Because either 7 gain or 10 loss must have occurred first in any tumor that has both copy number aberrations (CNAs), the fraction of tumors with both CNAs) at time $i$ is the sum of the fraction of tumors for which 7 changed first and the fraction of tumors for which 10 changed first.

$$\begin{aligned} P(\mathrm{gain7}_i, \mathrm{del10}_i) &= P(\mathrm{gain7}_i \mid \mathrm{del10}_{i-1})P(\mathrm{del10}_{i-1}) \\ &+P(\mathrm{del10}_i \mid \mathrm{gain7_{i-1}})P(\mathrm{gain7_{i-1}}) \end{aligned} \tag{1}$$

Care must be taken in interpreting this equation. If neither event has taken place, then the time point $i-1$ is undefined and probabilities such as $P(\mathrm{del10}_{i-1})$ are uninterpretable. Nonetheless, equation (1) holds formally if all probabilities are conditional on our stated model of $\mathrm{gain7_{i-1}}, \mathrm{gain7_I}$, $\mathrm{del10_{i-1}}$and $\mathrm{del10}_i$. One may write this condition as the following formal statement, where the standard symbols ∧, ⊕, ¬, and → to indicate logical "and", "xor'', "not'', and "implies" respectively.

$$(\mathrm{gain7_{i-1}} \oplus \mathrm{del10}_{i-1}) \wedge (\mathrm{gain7_{i-1}} \rightarrow \mathrm{gain7}_i) \wedge (\mathrm{del10}_{i-1} \rightarrow \mathrm{del10}_i) \tag{2}$$



2

We will not write this expression in our equations and inequalities, but rather assume informally that all probabilities are conditional on (2).

We wish to determine whether the following inequality holds.

$$\frac{1}{2}P(\mathrm{gain7}_i, \mathrm{del10}_i) < P(\,\mathrm{gain7}_i \mid \mathrm{del10}_{i-1}\,)P(\mathrm{del10}_{i-1}) \tag{3}$$

That is, whether most (greater than half) of the co-occurring mutations in the Progenetix set arise from loss of $10$ followed by gain of 7. Equation 3 does not indicate that we expect the left and right sides of the inequality to be nearly equal. In fact, one wants the two sides to be quite different to provide statistical evidence that the inequality does not just hold by chance.

The inequality (3) is inconveniently expressed in terms of time point $i-1$, whereas all observations were made at time point $i$. The time point of the conditional probabilities may be reversed using Bayes' Theorem write (3) equivalently as the following.

$$\frac{1}{2}P(\mathrm{gain7}_i, \mathrm{del10}_i) < P(\mathrm{del10}_{i-1} \mid \mathrm{gain7}_i)P(\mathrm{gain7}_i)$$

But it holds that:

$$\begin{aligned} P(\mathrm{del10}_{i-1} \mid \mathrm{gain7}_i)P(\mathrm{gain7}_i) &= P(\,\mathrm{del10}_i, \mathrm{del10}_{i-1} \mid \mathrm{gain7}_i\,)P(\mathrm{gain7}_i) \\ &= P(\,\mathrm{del10}_{i-1} \mid \mathrm{gain7}_i, \mathrm{del10}_i\,)P(\mathrm{gain7}_i, \mathrm{del10}_i) \end{aligned}$$

Because $\mathrm{del10}_{i-1}$implies $\mathrm{del10}_i$, (3) may be rewritten as

$$\frac{1}{2}P(\mathrm{gain7}_i, \mathrm{del10}_i) < P(\,\mathrm{del10}_{i-1} \mid \mathrm{gain7}_i, \mathrm{del10}_i\,)P(\mathrm{gain7}_i, \mathrm{del10}_i)$$

Cancelling $P(\mathrm{gain7}_i, \mathrm{del10}_i)$, which cannot be zero, we obtain the following inequality.

$$\frac{1}{2} < P(\,\mathrm{del10}_{i-1} \mid \mathrm{gain7}_i, \mathrm{del10}_i\,) = \frac{P(\mathrm{gain7}_i, \mathrm{del10}_{i-1})}{P(\mathrm{gain7}_i, \mathrm{del10}_i)} \tag{4}$$

At this point, we must approximate the right-hand side of (4) using the available data.

The right-hand side of equation (4) is the unknown fraction of tumors for which there are two changes but loss of 10 occurs first. To gain intuition, consider an unrealistic hypothetical set of tumors for which $P(\text{ gain7}_i \mid \text{del10}_{i-1}) = P(\text{ del10}_{i-1} \mid \text{gain7}_i)$. In such a case,

$$\frac{P(\text{gain7}_i, \text{del10}_{i-1})}{P(\text{gain7}_i, \text{del10}_i)} \approx \frac{P(\neg\text{gain7}_i \wedge \text{del10}_i)}{P(\text{gain7}_i \oplus \text{del10}_i)} \tag{5}$$

where the right-hand side of (5) represents the fraction of tumors for which one change, rather than two changes, has occurred and that change is loss of 10. The two sides of (5) are approximately equal because tumors acquire a second change at the same rate, independently of which change occurs first. Thus, estimating the right-hand side of (5) using the Progenetix data counts, in our contrived example

$$\frac{P(\text{gain7}_i, \text{del10}_{i-1})}{P(\text{gain7}_i, \text{del10}_i)} \approx \frac{\text{count}(\neg\text{gain7}_i \wedge \text{del10}_i)}{\text{count}(\text{gain7}_i \oplus \text{del10}_i)} \tag{6}$$

But empirical, longitudinal, data from Körber et al.[1] and related data from Barthel et al.[2] support the hypothesis that $P(\text{ gain7}_i \mid \text{del10}_{i-1}) \geq P(\text{ del10}_i \mid \text{gain7}_{i-1})$. If gain of 7 occurs preferentially after loss of 10, then instead of approximate equality, the right-hand side of (6) is expected to be a lower bound because the pool of samples with only loss of 10 will be preferentially depleted as a 7 gain occurs. The test

$$\frac{1}{2} < \frac{\text{count}(\neg\text{gain7}_i \wedge \text{del10}_i)}{\text{count}(\text{gain7}_i \oplus \text{del10}_i)} \tag{7}$$

is therefore meaningful unless loss of 10 does occur preferentially after gain of 7.

Importantly, Progenetix does not supply longitudinal data that justify the less-than sign in (7); data from from Körber et al.[1] and Barthel et al.[2] must be used. Similarly, longitudinal data cannot supply the numerical counts required for the test of inequality (7); a collection of tumor data, such as Progenetix, is needed.

The numbers of brain cancer samples in the Progenetix data and having each of three mutually exclusive presence/absence of 7 gain and 10 loss are in the following table.

| | | |
|---|---|---|
| $\text{gain7}_i$ | $\text{del10}_i$ | 1214 |


4

| | | |
|---|---|---|
| $\neg\text{gain7}_i$ | $\text{del10}_i$ | 555 |
| $\text{gain7}_i$ | $\neg\text{del10}_i$ | 339 |
| Total satisfying condition (1) | | 2108 |

Using the data in this table, one may test

$$\frac{1}{2} < \frac{\text{count}(\neg\text{gain7}_i \wedge \text{del10}_i)}{\text{count}(\text{gain7}_i \oplus \text{del10}_i)} = \frac{555}{555+339} = \frac{555}{894} \approx 0.62 \quad (8)$$

The right-hand side follows a binomial distribution with 555 successes out of 894 trials and is different from 1/2 with a one-sided p-value of 2.50e-13.

The test corresponding to (6) for the opposite assumption – that at least half the samples are derived from a gain of 7 followed by a loss of 10 -- is the following.

$$\frac{1}{2} < \frac{\text{count}(\text{gain7}_i \wedge \neg\text{del10}_i)}{\text{count}(\text{gain7}_i \oplus \text{del10}_i)} = \frac{339}{555+339} = \frac{339}{894} \quad (9)$$

But 339/894 is significantly less than 1/2 and so a one-sided binomial test fails with an extreme p-value of approximately 1. Thus, on the assumption that loss of 10 does not happen preferentially after gain of 7, inequalities (6) and (7) are consistent with more than half of the Progenetix samples arising from a loss of 10 followed by a gain of 7, with an estimated fraction that is at least 0.62. Given the number of samples, 0.62 is significantly greater than 0.5 with a p-value of 2.50e-13.

**2. Preferential loss of 10 before gain of 7 is sufficient to explain preference for the co-occurrence**

One key question our study addresses is why is it plausible the joint probability of a 7 gain and a 10 loss is much larger than the product of the probabilities of 7 gain and 10 loss as single events? We define this difference as the 'excess probability' of 7 gain and 10 loss. Since we justified above



5

that the more common order is 10 loss first and 7 gain second, we expect that some of the excess probability is explained by tumors in which the two events occurred in the more common order and that is why we investigated the reasons why 7 gain following 10 loss may confer a fitness benefit. In Note 2, we address the probabilistic question: are tumors in which 10 loss is followed by 7 gain sufficient to explain all the excess probability? If that is the case, then we do not need to investigate whether the 'minority' occurrence of 10 loss after 7 gain confers any fitness benefit.

Let $x$ be the fraction of tumors having both a $10$ loss and $7$ gain for which the $10$ loss happened first, assuming we could know the order of events. By equation (1), since the events are exclusive, $1-x$ is the fraction of minority tumors for which 7 gain happens before 10 loss. If the two possible orders were equally likely, then $x=0.5$, and $1-x=0.5$. Let us first investigate excess probability with an equally likely sequence of events. Let $P(X)$ indicate the observed probability of event $X$, without the implicit conditioning of the previous section. Let $P_{M0.5}(X)$ be the probability of event $X$ based on the simplified model that the two events are occur in either order with equal probabilities.

$$P_{M0.5}(X) = P(7\text{ gain})P(10\text{ loss}) \tag{10}$$

All observations are at time $i$, so we drop the subscript $i$ in the notation. The excess probability that is attributable to the minority (10 loss comes second) tumors

$$\begin{aligned} M(0.5) &= P(7\text{ gain}, 10\text{ loss}) - P_{M0.5}(7\text{ gain}, 10\text{ loss}) \\ &= P(7\text{ gain}, 10\text{ loss}) - P(7\text{ gain})P(10\text{ loss}) \end{aligned}$$

From the Progenetix counts in **Fig. 1B**, it follows that:

$$P(7\text{ gain}) = \frac{1214+339}{2813} \approx 0.552$$

$$P(10\text{ loss}) = \frac{1214+555}{2813} \approx 0.629$$

$$P(7\text{ gain}, 10\text{ loss}) = \frac{1214}{2813} \approx 0.432$$

And therefore, that the excess joint probability attributable to the minority tumors, if $x=0.5$ would be

$$M(0.5) = 0.432 - (0.552 \text{ x } 0.629) \approx 0.085$$

Thus, if $x = 0.5$ , which corresponds to an equally likely order of the two aneuploidies, the observed preferred order tumors with 10 loss first and 7 gain second are not sufficient to explain the observed probability of co-occurrence because the excess probability exceeds 0.

Let us now formulate a model for $x \neq 0.5$. In this formulation, we imagine that the tumors with 7 gain after 10 loss are conceptually partitioned into two categories: tumors in which the second event is independent of the first and others in which the second event is deterministic. This is a probabilistic technique called a "mixture model" to attribute how much more likely 7 gain is to occur after 10 loss than in the absence of 10 loss. The difference between the probabilities, $x - (1 - x) = 2x - 1$ is effectively the proportion of tumors in which 7 gain occurs 'deterministically' after 10 loss. In the previous section we established that it is significantly likely that $x > 0.5$, and so we can assume that $x$ lies in the interval $[0.5, 1]$ and therefore that $2x - 1$ lies in [0,1]. Our model takes the following form.

$$P_{Mx}(7\text{ gain}, 10\text{ loss}) \approx (2 - 2x)P(7\text{ gain})P(10\text{ loss}) + (2x - 1)\ P(\text{ 7 gain} \mid 10\ loss\,)$$

Note that $2 - 2x + 2x - 1 = 1$ as it must in a mixture model, that $2 - 2x$ lies in $[0,1]$, and that the model reduces to (8) in the case where $x = 0.5$.

We get the following weighted excess probability attributable to the minority tumors.

$$M(x) \approx 0.432 - 0.348(2 - 2\text{x}) - 0.687(2\text{x} - 1)$$
$$\approx 0.424 - 0.678\text{x} \qquad (11)$$

The equality is approximate because there are minor discrepancies created by rounding the third significant digit. In the above equation (11), 0.348 = P(7 gain) P(10 loss) = 0.552 x 0.659. Also, 0.687 = P(7 gain | 10 loss) = P(7 gain and 10 loss) / P(10 loss) = 0.432/0.629. The linear equation (11) is plotted in **Fig. S1** in a manner that reduces our question to a root finding problem; the x-axis ranges unconventionally from (0.5,1] because, as explained above, $x > 0.5$. We want to find the unique value of *x* at which the model has exactly zero excess probability (remaining to be explained by the tumors with 7 gain first and 10 loss second). We then wish to evaluate if this $x$ is consistent with the empirical data of Körber et al.[1]. This consistency check is not circular reasoning because those additional data are not in Progenetix and were not used to derive the model.

Equation (11) has its single root at $x$ = 0.424/0.678 ≈ 0.625, which approximately corresponds to 5/8 of the data having loss of 10 followed by gain of 7. The data in Figure 4A of Körber et al.[1] suggest an empirical estimate of $x \geq 0.5 + \frac{2}{21} = 0.595$, which is not significantly different from 0.625 by a binomial test. Thus, we conclude that all the excess probability of co-occurrence can be explained by tumors in which 10 loss occurs first and 7 loss occurs second. This indicates that the opposite order of events in which 7 gain occurs first and 10 loss occurs second can be treated as occurring by random chance meaning that 10 loss is not more likely to occur after 7 gain occurred. The 10 loss occurring second may ultimate be explained by some yet-to-be-determined mechanism, but we do not need to explore whether 10 loss after 7 gain confers fitness benefits (deriving from some interaction between the two chromosomes).

**3. Pairwise pathway enrichment**

To identify the pathway pairs enriched by these DU-SR gene pairs, we carried out a pairwise pathway enrichment analysis among the vulnerable genes located on chromosome 10 and their rescuer genes on chromosome 7, using 50 Cancer Hallmark pathways (MSigDB's hallmark genesets[3]). The DU-SR network generated from TCGA GBM+LGG data. We employed a one-sided Fisher exact test to examine whether the gene pairs (comprising vulnerable and rescuer genes) are statistically enriched in specific pairs of pathways. This aims to identify pathways whose dysfunction can be mitigated by the activity of rescuer genes in other pathways. FDR correction was applied to adjust for multiple hypothesis testing.

We observed that vulnerable genes on chromosome 10 that are involved in glycolysis, heme metabolism, and hypoxia are rescued by chromosome 7 genes involved in pathways including notch signaling, Wnt beta-catenin signaling, and epithelial-mesenchymal transition. The


8

full list of such enriched pathway pairs is summarized in **Fig. S4**, **Table S5**. For **Fig. S4A**, we focus solely on the DU-SR network involving vulnerable genes on chromosome 10 and rescuer genes on chromosome 7, using all the genes on these chromosomes as the universal background set for enrichment. For **Fig. S4B**, the DU-SR network includes vulnerable genes on chromosome 10 and rescuer genes from any chromosome, with the universal background set extending to all genes on chromosome 10 and any chromosome, respectively.




9

**SUPPLEMENTARY FIGURES**

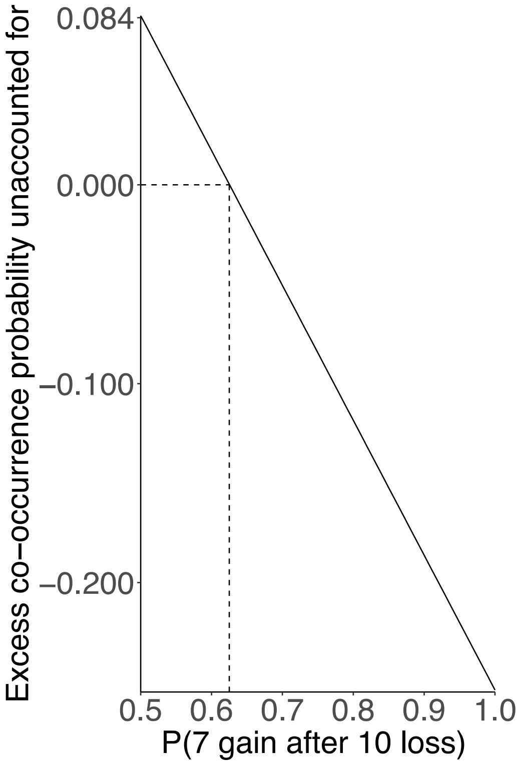


***Figure S1:*** *A mathematical model computing P(7 gain after 10 loss) in GBM from the Progenetix data. The dotted line is the value of P(7 gain after 10 loss) when the solution to the equation is 0. P(x) stands for probability of event x.*

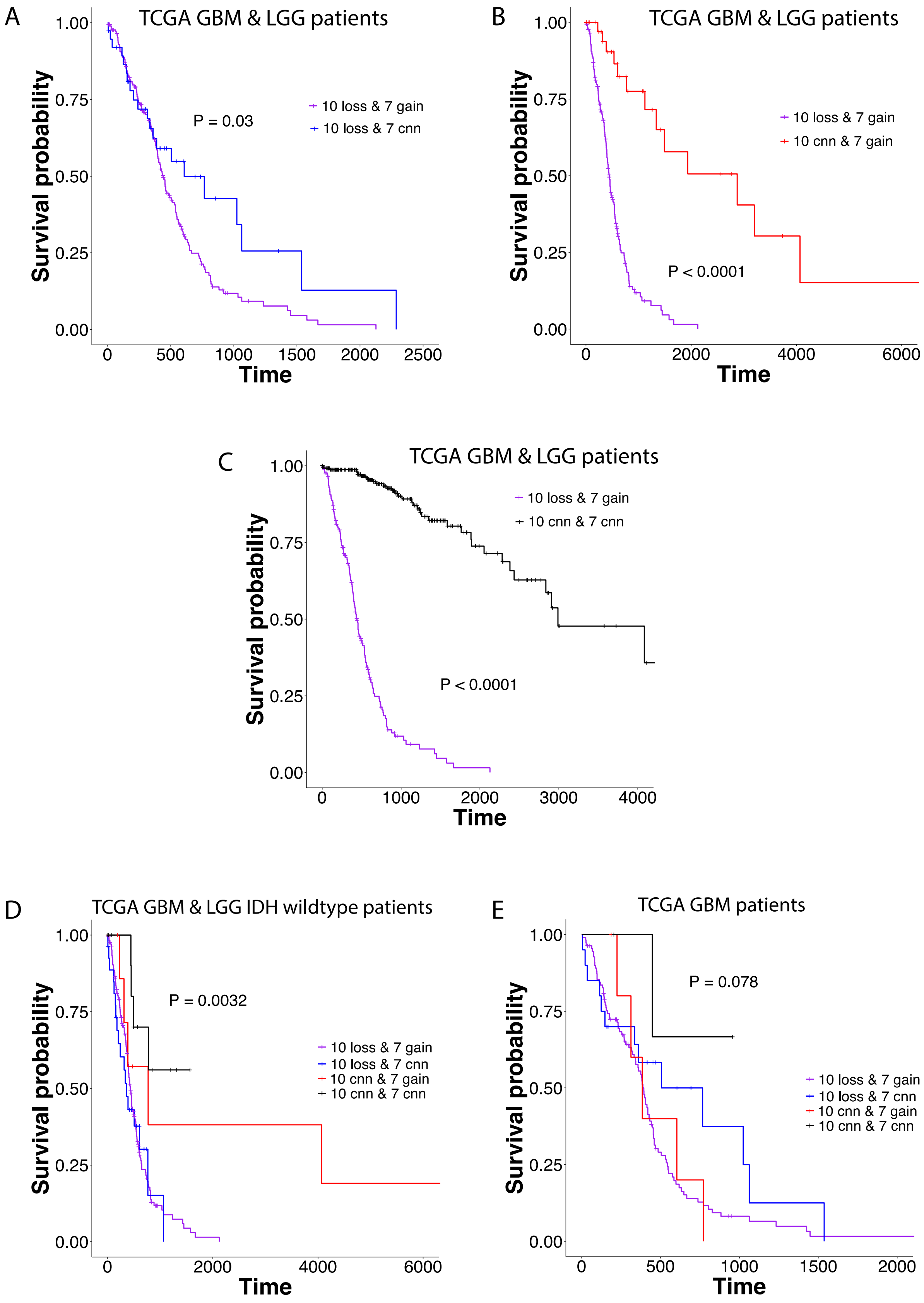
A
TCGA GBM & LGG patients
P = 0.03
10 loss & 7 gain
10 loss & 7 cnn
Survival probability
Time
B
TCGA GBM & LGG patients
P < 0.0001
10 loss & 7 gain
10 cnn & 7 gain
Survival probability
Time
C
TCGA GBM & LGG patients
P < 0.0001
10 loss & 7 gain
10 cnn & 7 cnn
Survival probability
Time
D
TCGA GBM & LGG IDH wildtype patients
P = 0.0032
10 loss & 7 gain
10 loss & 7 cnn
10 cnn & 7 gain
10 cnn & 7 cnn
Survival probability
Time
E
TCGA GBM patients
P = 0.078
10 loss & 7 gain
10 loss & 7 cnn
10 cnn & 7 gain
10 cnn & 7 cnn
Survival probability
Time



11

***Figure S2:*** *Survival analysis (Kaplan-Meier curves) in TCGA GBM/LGG patients with the following occurrences: 10 cnn & 7 cnn, 10 loss & 7 cnn, 10 cnn & 7 gain, 10 loss & 7 gain. Log-rank test p-value (P) is shown. Survival analysis is done using TCGA GBM and LGG data in pairwise manner comparing 10 loss & 7 gain patient samples with (n=174):* ***(A)*** *10 loss and 7 cnn (n=38);* ***(B)*** *10 cnn and 7 gain (n=40);* ***(C)*** *10 cnn and 7 cnn (n=259). Survival analysis is done only for* ***(D)*** *IDH wildtype TCGA GBM/LGG patients (n = 15 for 10 cnn & 7 cnn, n = 27 for 10 loss & 7 cnn, n = 9 for 10 cnn & 7 gain, n = 168 for 10 loss & 7 gain);* ***(E)*** *TCGA GBM patients (no LGG patients) (n = 4 for 10 cnn & 7 cnn, n = 20 for 10 loss & 7 cnn, n = 6 for 10 cnn & 7 gain, n = 111 for 10 loss & 7 gain). Keywords – GBM: Glioblastoma multiforme, LGG: brain lower grade glioma; 10 cnn or 7 cnn: samples with copy number neutral state for chromosomes 10 or 7, respectively; '&' implies 'and'; 10 loss or 7 gain: samples with chromosome 10 lost or chromosome 7 gained, respectively.*



12

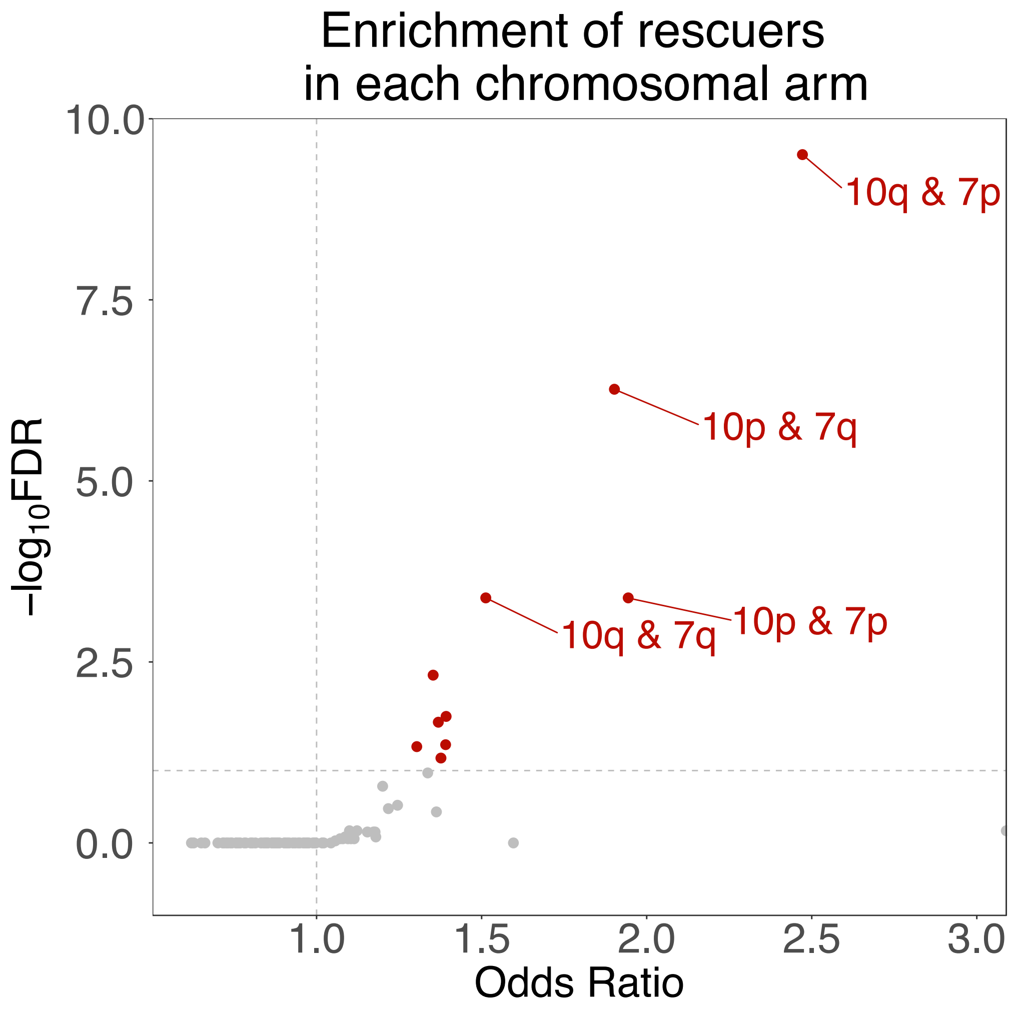


***Figure S3:*** *We repeated the DU SR analysis on TCGA GBM and LGG samples by explicity removing one of the steps of INCISOR which looks for positive selection of gene pairs with high expression or copy-number variation. Volcano plot showing the overlap enrichment of rescuer genes with the genes in each chromosomal arm (Fisher exact test). The dashed horizontal line is FDR = 0.1, and the dashed vertical line is at odds ratio = 1. The rescuer genes are identified for all the (vulnerable) genes on a specific arm and the tested for overlap enrichment across all chromosomal arms. Rescuers of vulnerable genes in 10p or 10q are more enriched in chromosomes 7p or 7q. Half-dot implies odds ratio = Inf.*

A

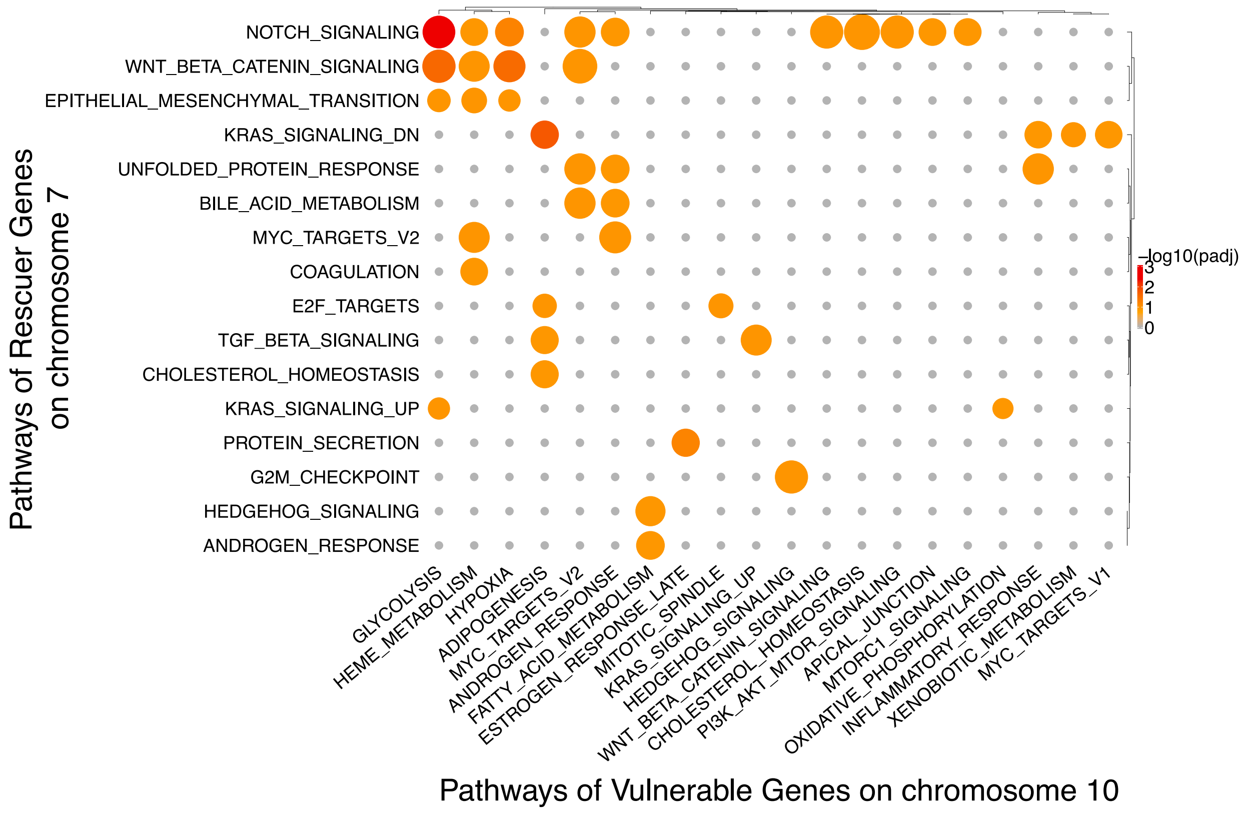

NOTCH_SIGNALING
WNT_BETA_CATENIN_SIGNALING
EPITHELIAL_MESENCHYMAL_TRANSITION
KRAS_SIGNALING_DN
UNFOLDED_PROTEIN_RESPONSE
BILE_ACID_METABOLISM
MYC_TARGETS_V2
COAGULATION
E2F_TARGETS
TGF_BETA_SIGNALING
CHOLESTEROL_HOMEOSTASIS
KRAS_SIGNALING_UP
PROTEIN_SECRETION
G2M_CHECKPOINT
HEDGEHOG_SIGNALING
ANDROGEN_RESPONSE
Pathways of Rescuer Genes on chromosome 7
GLYCOLYSIS
HEME_METABOLISM
HYPOXIA
ADIPOGENESIS
MYC_TARGETS_V2
ANDROGEN_RESPONSE
FATTY_ACID_METABOLISM
ESTROGEN_RESPONSE_LATE
MITOTIC_SPINDLE
KRAS_SIGNALING_UP
HEDGEHOG_SIGNALING
WNT_BETA_CATENIN_SIGNALING
CHOLESTEROL_HOMEOSTASIS
PI3K_AKT_MTOR_SIGNALING
APICAL_JUNCTION
MTORC1_SIGNALING
OXIDATIVE_PHOSPHORYLATION
INFLAMMATORY_RESPONSE
XENOBIOTIC_METABOLISM
MYC_TARGETS_V1
Pathways of Vulnerable Genes on chromosome 10
-log10(padj)
3
2
1
0




14

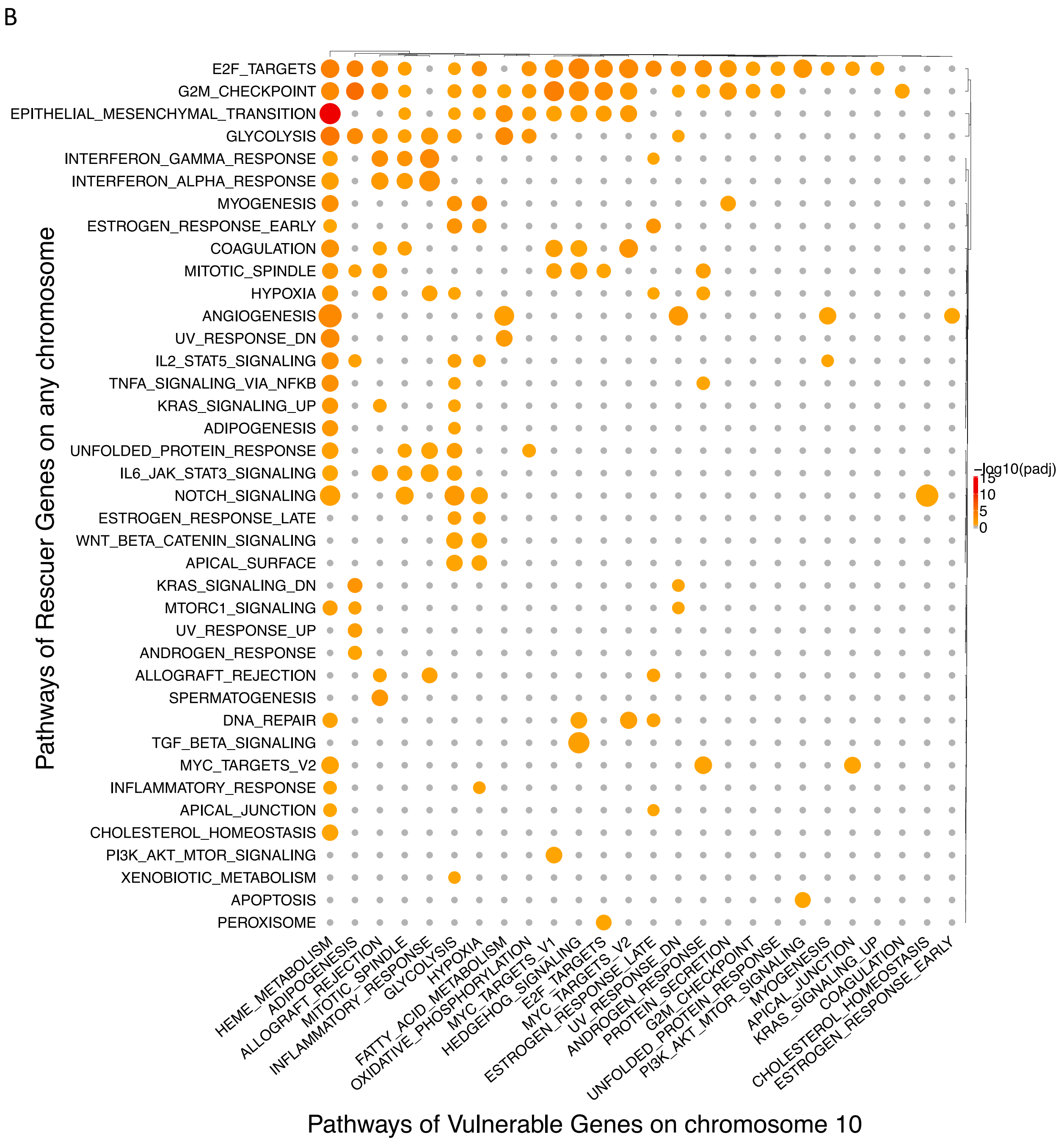


***Figure S4:*** *Pairwise pathway enrichment analysis for the DU-SR network derived from GBM+LGG data for the vulnerable genes in chromosome 10 and rescuer genes in:* ***(A)*** *chromosome 7 and* ***(B)*** *any chromosome. Statistical test used is Fisher exact test (P < 0.05, adjusted-P/FDR < 0.2). The established set of 50 Cancer Hallmark pathways was used for this analysis (MSigDB's hallmark genesets[3]). Keywords: 'padj' implies 'adjusted p-value'.*



15

*Pairwise pathway enrichment analysis for the DU-SR network derived from GBM+LGG data for the vulnerable genes in chromosome 10 and rescuer genes in any chromosome (Fisher exact test, P < 0.05, adjusted-P/FDR < 0.2).*



16

A

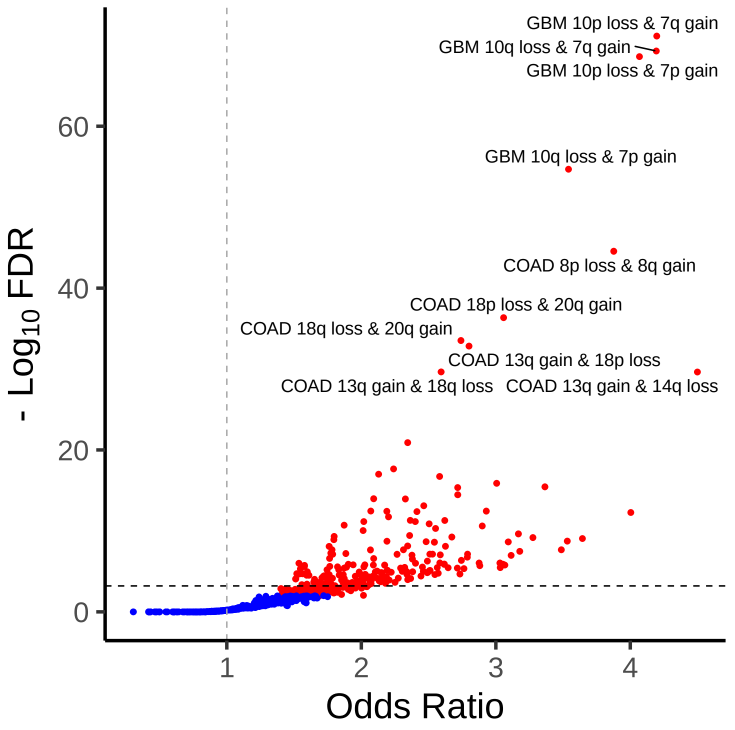
GBM 10p loss & 7q gain
GBM 10q loss & 7q gain
GBM 10p loss & 7p gain
GBM 10q loss & 7p gain
COAD 8p loss & 8q gain
COAD 18p loss & 20q gain
COAD 18q loss & 20q gain
COAD 13q gain & 18p loss
COAD 13q gain & 18q loss
COAD 13q gain & 14q loss
60
40
20
0
- Log10 FDR
1
2
3
4
Odds Ratio

B

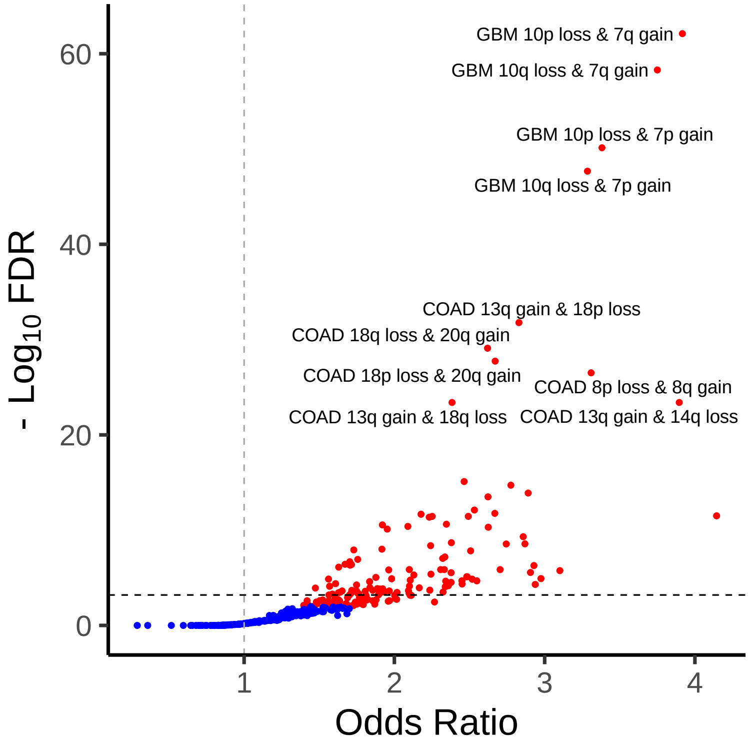
GBM 10p loss & 7q gain
GBM 10q loss & 7q gain
GBM 10p loss & 7p gain
GBM 10q loss & 7p gain
COAD 13q gain & 18p loss
COAD 18q loss & 20q gain
COAD 18p loss & 20q gain
COAD 8p loss & 8q gain
COAD 13q gain & 18q loss
COAD 13q gain & 14q loss
60
40
20
0
- Log10 FDR
1
2
3
4
Odds Ratio



17

C

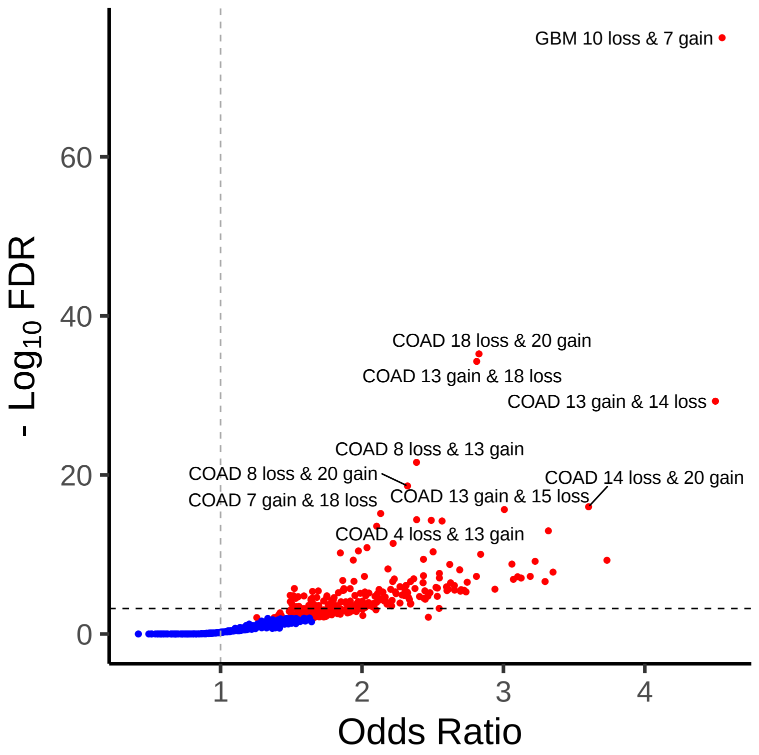


***Figure S5:** Plot charting the landscape of chromosomal arm loss-gain enrichments across various cancer types from patient tumors in Progenetix dataset. (A) Chromosomal arm gain/loss events at a threshold of 60% of the arm called gained or lost. (B) Chromosomal arm gain/loss events at a threshold of 80% of the arm called gained or lost. (C) Chromosomal gain/loss events when the loss or gain of a either arm is considered loss or gain of the chromosome.*



18
